# Control of dynamic cell behaviors during angiogenesis and anastomosis by Rasip 1

**DOI:** 10.1101/2020.08.27.269977

**Authors:** Minkyoung Lee, Charles Betz, Ilkka Paatero, Niels Schellinx, Jianmin Yin, Christopher William Wilson, Weilan Ye, Markus Affolter, Heinz-Georg Belting

## Abstract

Organ morphogenesis is driven by a wealth of tightly orchestrated cellular behaviors, which ensure proper organ assembly and function. Many of these cell activities involve cell-cell interactions and remodeling of the F-actin cytoskeleton. Here, we analyze the requirement for Rasip1 (Ras-interacting protein 1), an endothelial-specific regulator of junctional dynamics, during blood vessel formation. Phenotype analysis of *rasip1* mutants in zebrafish embryos reveal distinct requirements for Rasip1 during sprouting angiogenesis, vascular anastomosis and lumen formation. During angiogenic sprouting, Rasip1 is required for efficient cell pairing, which is essential for multicellular tube formation. High-resolution time-lapse analyses show that these cell pairing defects are caused by a destabilization of tricellular junctions suggesting that tri-cellular junctions may serve as a counterfort to tether sprouting endothelial cells during morphogenetic cell rearrangements. During anastomosis, Rasip1 is required to establish a stable apical membrane compartment; *rasip1* mutants display ectopic, reticulated junctions and the apical compartment is frequently collapsed. Loss of Ccm1 and Heg1 function leads to junctional defects similar to those seen in *rasip1* mutants. Analysis of *radil-b* single and *rasip1/radil-b* double mutants reveal distinct and overlapping functions of both proteins. While Rasip1 and Radil-b have similar functions during angiogenic sprouting, the junction formation during anastomosis may primarily depend on Rasip1.

## Introduction

The cardiovascular system is the first organ to become functional during embryonic development. The generation of vascular networks is essential for developmental patterning, growth and survival of the vertebrate embryo. As the embryo grows, the vasculature adjusts to the increasing demand of nutrients and oxygen by an expansion of the vasculature tree via sprouting angiogenesis, vascular remodeling and adaptation of blood vessel diameter. Vascular morphogenesis is driven by a wealth of dynamic cellular behaviors, which are regulated by molecular as well as physical cues, and are characterized by an extraordinary plasticity (Adams and Alitalo, 2007; Baeyens et al., 2016; Duran et al., 2017). At the cellular level, blood vessel morphogenesis and remodeling are accomplished by endothelial cell behaviors including cell migration, cell rearrangement and cell shape changes (Betz et al., 2016). This repertoire of dynamic behaviors allows endothelial cells to rapidly respond to different contextual cues, for example during angiogenic sprouting, anastomosis, pruning, diapedesis or regeneration.

Previous studies have shown that vascular tube formation requires extensive and diverse cell shape changes and that these changes can be driven by junctional remodeling as well as dynamic regulation of the cortical actin cytoskeleton (Gebala et al., 2016; Paatero et al., 2018; Phng et al., 2015; Sauteur et al., 2014). Junctional remodeling is essential for cell rearrangements, which drive multicellular tube formation. Enlargement of the luminal space, on the other hand, requires apical membrane invagination (Barry et al., 2016; StriliC et al., 2009). During anastomosis, the apical membrane can invaginate through the entire cell leading to the formation of a unicellular tube (Lenard et al., 2013).

Junctional remodeling and membrane invagination rely on the dynamic regulation of the F-actin cytoskeleton at the endothelial cell junction and apical cortex, respectively. Small GTPases of the Rho family, including Cdc42, Rac1 and RhoA are essential regulators of F-actin dynamics and have been shown to play critical roles during blood vessel formation *in vitro* and *in vivo* (reviewed by (Barlow and Cleaver, 2019). In the vasculature, these GTPases are partially regulated by the adaptor protein Rasip1. Rasip1 has been shown to promote Rac1 and Cdc42 activity, whereas it inhibits RhoA activity by binding to the GTPase activating protein Arhgap29 (Barry et al., 2016; Xu et al., 2011). Ablation of Rasip1 in mice and knock-down of *rasip1* zebrafish cause severe vascular defects (Wilson et al., 2013; Xu et al., 2011). During vasculogenesis, Rasip1 is required for the lumenization of the dorsal aorta, in particular for the clearing of apical membrane compartments from junctional proteins and for the opening of the vascular lumen between endothelial cells (Barry et al., 2016). However, the role of Rasip1 during sprouting angiogenesis and anastomosis has not been studied in detail. To gain more insight into the cellular and molecular mechanisms of vascular tube formation during angiogenesis, we have generated loss-of-function alleles in the zebrafish *rasip1* gene and performed high-resolution time-lapse imaging to observe junctional dynamics during sprouting angiogenesis and anastomosis. Loss of *rasip1* causes multiple vascular defects, with respect to angiogenic sprouting, including defects in cell proliferation, junctional stability and lumen formation. Furthermore, analyses of *radil-b* and *rasip1;radilb* double mutants reveal partly redundant roles for the two proteins. Lastly, knock-down of *ccm1* and *heg1* phenocopies the apical junctional defects seen in *rasip1*, suggesting a functional interaction between the proteins during blood vessel formation.

## Results

### Loss of Rasip1 function causes broad vascular defects

To investigate the role of Rasip1 in vascular morphogenesis we employed CRISPR/Cas9 technology to generate several mutant alleles, namely *rasip1*^*ubs23*^, *rasip1*^*ubs24*^ and *rasip1*^*ubs28*^, respectively (S-Figure 1). The *rasip1*^*ubs28*^ allele comprises a deletion of about 35kb including the *rasip1* coding region from exon 3 to 16, encoding a severely truncated protein lacking the Ras-association, Forkhead-association and Dilute domains (Figure 1a). Since the truncated protein lacks all the conserved domains, we consider *rasip1*^*ubs28*^ to be a null-allele and focused our studies on the analysis of this mutant.

**Figure 1:**
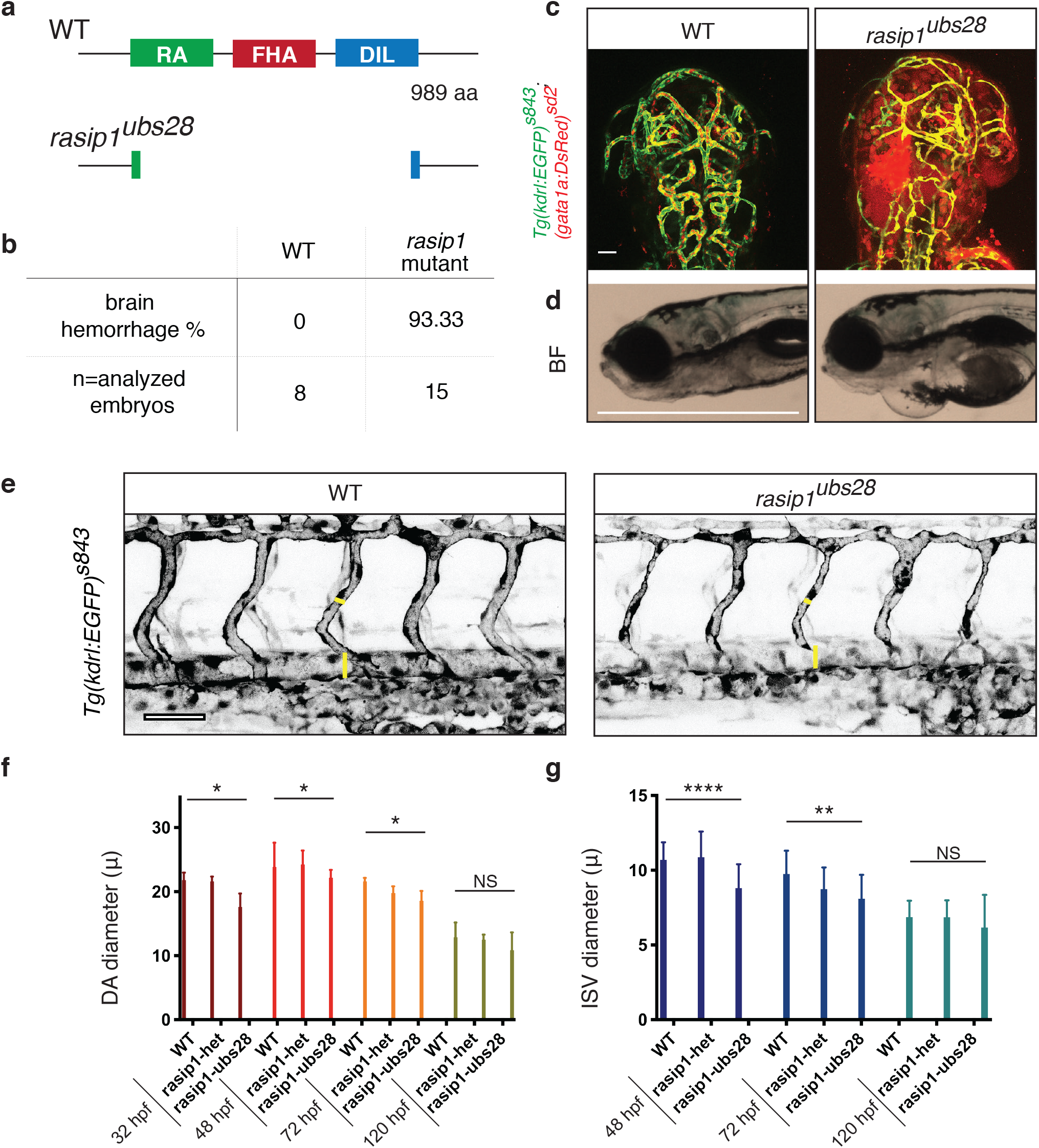
Vascular defects in zebrafish *rasip1* mutants. **(a)** Conserved Rasip1 protein domains. RA: Ras association domain. FHA forkhead-association domain. DIL: dilute domain. The *rasip1*^*ubs28*^ mutant allele consists of a 35kb deletion comprising all three domains. **(b)** quantification of cranial hemorrhages in *rasip1*^*ubs28*^ mutant embryos. p<0.0001 (Fisher’s exact test). **(c)** confocal images of cranial hemorrhages in rasip1^***ubs28***^ embryos (72 hpf). Blood cells are visualized by gata1:dsRed expression. Scale bar, 50 μm. **(d)** bright field image (BF) of wild-type and *rasip1*^*ubs28*^ embryos showing pericardial edema. Scale bar, 2 mm. **(e)** confocal images of wild-type and *rasip1*^*ubs28*^ mutants. Mutants show narrower DA and irregular ISV diameters. Scale bar, 50 μm. **(f)** quantification of DA diameters (µm) during embryonic development (32 to 120 hpf). Mann-Whitney test and error bars indicate standard deviation; significance (ns=not significant, *p < 0.1). (WT 32, 48, 72, 120 hpf: *n*=4, 4, 9, 13 embryos; *rasip1*^*ubs28/+*^ *n*=5, 12, 19, 33; *rasip1*^*ubs28*^ *n*=6, 10, 9, 10). **(g)** quantification of ISV diameters (µm) during embryonic development (32 to 120 hpf). Embryos were analyzed by unpaired two-tailed Mann-Whitney test and error bars indicate standard deviation; significance (ns=not significant, *p < 0.1, **p < 0.01, ****p < 0.0001). (WT 48, 72, 120 hpf: *n*=37, 94, 70 ISVs; *rasip1*^*ubs28/+*^ *n*=104, 52, 198; *rasip1*^*ubs28*^ *n*=67, 38, 65).

Homozygous *rasip1* mutants displayed hemorrhages and vascular instability in the cranial vasculature at 3 days of development (Figure 1b,c). Furthermore, we observed transient pericardial edema between 3 and 5 dpf (Figure 1d), which usually disappeared during larval development. In the trunk, *rasip1* mutants showed reduced blood flow, which correlated with irregular and generally reduced diameter of the dorsal aorta (DA) and intersegmental vessels (ISVs) (Figure 1e-g). These differences were transient and by 5 dpf the average vessel diameter had recovered to normal size. Despite these severe defects, homozygous *rasip1* mutants recovered and about 30% of them developed to fertility.

### Reduced motility and proliferation during angiogenic sprouting in *rasip1* mutants

To find out whether the loss of *rasip1* may affect dynamic cell behaviors, we performed time-lapse analyses, concentrating on the developing ISVs. ISVs sprouts emerge from the DA at about 22-24 hpf and extend towards the dorsal side of the embryo. In wild-type embryos, they reach the dorsal part of the neural tube by 28 hpf and initiate formation of the dorsal longitudinal anastomotic vessel (DLAV) (Lawson and Weinstein, 2002). In our time-lapse movies we observed that by 30 hpf almost all sprouts in the trunk region had completed dorsal sprouting and engaged in contact with neighboring sprouts (S-Figure 2a, b). In contrast, ISV sprouting in *rasip1* mutants appeared sluggish and 25% of sprouts were incomplete by 30 hpf (S-Figure 2a,b). To see whether stunted outgrowth was reflected by a difference in the number of endothelial cells contributing to ISV sprouts, we counted nuclei within each sprout at 30 hpf (S-Figure 2c). In wild-type and *rasip1* mutants, the number ISV nuclei was quite variable ranging between one and five. However, we observed a clear enrichment of ISVs containing one or two nuclei in mutants when compared to wild-type, which contained three to four nuclei. To differentiate whether this diminished cell number was caused either by reduced recruitment or by proliferation defects of endothelial cells, we tracked endothelial cell nuclei during ISV formation (S-Figure 2d-g). In wild-type siblings, we observed that two cells migrated from the DA into the sprout, undergoing one round of division each thus giving rise to an ISV consisting of four cells (S-Figure 2e); ISVs comprising three cells were usually formed by two migrating cells and a single cell division (S-Figure 2f). In *rasip1* mutants we rarely observed cell divisions within the sprouts (S-Figure 2d,g). Instead, most cells in the sprout originated from the DA and occasionally we observed three cells migrating into the sprout (Figure 2f). Hence, these results show that proliferation in sprouting endothelial cells is reduced in *rasip1* mutants and suggest that paucity of cell number may partially be compensated by the recruitment of additional cells into the sprout.

**Figure 2:**
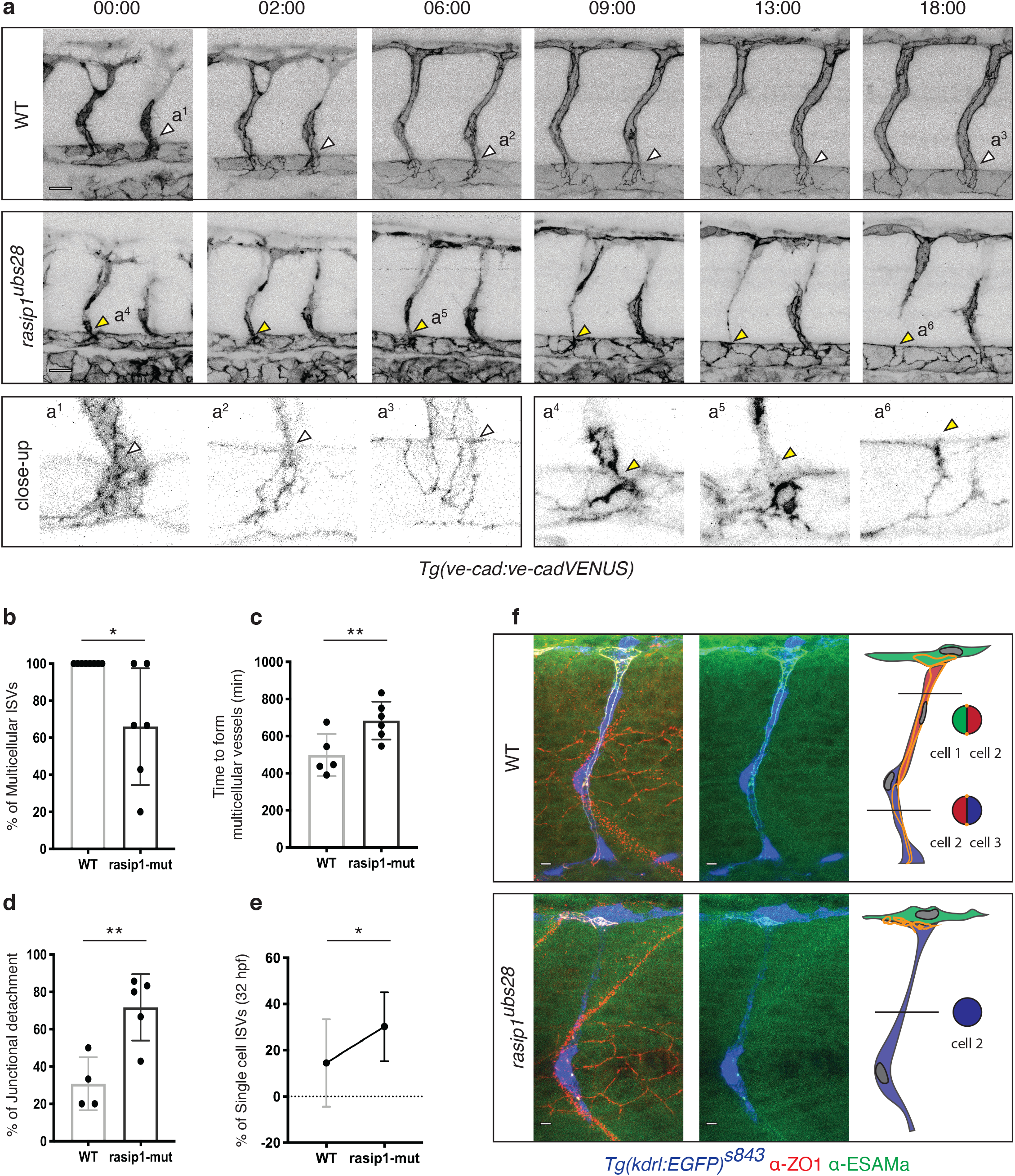
Formation of multicellular vessels is impaired in *rasip1* mutants. **(a)** Still pictures of time-lapse movies (s-movies 7 and 8) showing endothelial cell junctions (Cdh5-Venus) in wild-type and *rasip1*^*ubs28*^ embryos. White arrowheads show maintained junctional in wild-type ISV sprouts. Yellow arrowheads indicate junctional detachment in mutant embryos. Scale bars, 20 μm. Bottom row: closeups of showing junctional detachment in a^4^ and a^5^. **(b-e)** Quantification of junctional and cellular configuration during ISV formation in wild-type and *rasip1*^*ubs28*^ mutant embryos. **(b)** Percentage multicellular tubes at 48 hpf (wt *n*=8, mut *n*=6). **(c)** Speed of multicellular tube formation (wt *n*=5, mut *n*=6). **(d)** Percentage of ISV with multicellular configuration per embryo (WT *n*=4, mut *n*=5). **(e)** Percentage of single cell ISVs at 32 hpf (WT *n*=8, mut *n*=8). Quantifications were done by counting ISVs showing the respective phenotypes, averaged by total ISVs analysed per embryo. **(f)** Immunofluorescence of Zo-1 and Esama in *Tg(kdrl:EGFP)*^*s843*^ at 32 hpf. Schematic drawings on the right show the different cellular configurations of multicellular (wt) and unicellular (*rasip1* mutant) ISVs. Scale bars, 5 μm. The data was analyzed by unpaired two-tailed Mann-Whitney test and error bars indicate standard deviation; significance (*p < 0.1, **p < 0.01).

### Instability of tricellular junctions inhibits formation of multicellular tubes

We next examined whether loss of *rasip1* affects angiogenic tube formation. In wild-type embryos, the multicellular architecture of ISVs tubes is established via concerted migratory and proliferative activities of endothelial cells. More specifically, the multicellular configuration is driven by junctional rearrangements along the vessel axis, which leads to effective cell pairing and thus multicellularity.

Since multicellular tubes are characterized and can be recognized by continuous junctions along the blood vessel axis (Blum et al., 2008), we used a *ve-cad:ve-cad-Venus* reporter (Lagendijk et al., 2017) to follow the dynamics of endothelial junctions during ISV formation. In wild-type siblings, cell junctions elongated and spanned the entire extend of the ISV giving rise to multicellular tubes by 48 hpf (Figure 2a,b,f). Notably, adherens junctions maintained the continuity of the ISVs with the DA, where they formed vertices or tri-cellular junctions (Figure 2a, white arrowheads). *rasip1* mutants showed a clear delay in multicellular tube formation (Figure 2c) and at 48 hpf – on average – about 40% of ISVs had not achieved a multicellular configuration (Figure 2b). Moreover, time-lapse analysis of VE-cad-Venus showed defects in junctional development (Figure 2a). Specifically, at the ventral base of the sprout, junctions that were normally tethered to the DA in wild-type embryos, lost this attachment and the junctional ring was “released” in mutant embryos (Figure 2a, yellow arrowheads,d). This detachment resulted in one of the stalk cells moving up into the DLAV leaving a single cell spanning the distance between the DA and the DLAV (Figure 2a,e,f). These results indicate that junctional interconnections at the base of the sprout are critical for cell intercalation to occur during multicellular tube formation. Thus, the loss of these connections in *rasip1* mutant prevents cell pairing and results in unicellular ISVs and consequently a defect in the cord hollowing process underlying multicellular tube formation.

### Defects in junctional dynamics during blood vessel anastomosis

The growth and interconnection of vascular networks requires angiogenic sprouting as well as the interconnection of all sprout by the process of anastomosis. At the cellular level, anastomosis occurs in a quite stereotyped fashion (reviewed by (Betz et al., 2016). Neighboring sprouts initiate contact via tip cell filopodia and form a junctional ring which surrounds apically polarized membrane. Junctional ring and apical membrane formation in both of the contacting cells leads to the formation of a luminal pocket, which is later connected to the lumen of the nascent vessel. The process of anastomosis in zebrafish serves as a paradigm to study the cell biology of blood vessel formation and includes processes such as apical polarization, junctional rearrangements and lumen formation, which occur within a 4 to 6 hours (Herwig et al., 2011). To assess the role of Rasip1 during anastomosis, we compared the dynamics of junctional reporters such as VE-cad-Venus (Figure 3a) and Pecam-EGFP (S-Figure 3) in wild-type and mutant embryos. Time-lapse analyses revealed two different defects in *rasip1* mutants during junctional ring formation. In about 53% of anastomosis events (Figure 3a) we observed ectopic accumulation of VE-cadherin-Venus or Pecam-EGFP (S-Figure 3) within the junctional ring, revealing a defect in relocating these junctional proteins from the apical compartment to cell junctions. Alternatively, in about 34% of cases, the anastomotic ring (Figure 3c,e,f) elongated along the blood vessel axis but failed to maintain a lateral axis, leading to a collapsed junctional ring.

**Figure 3:**
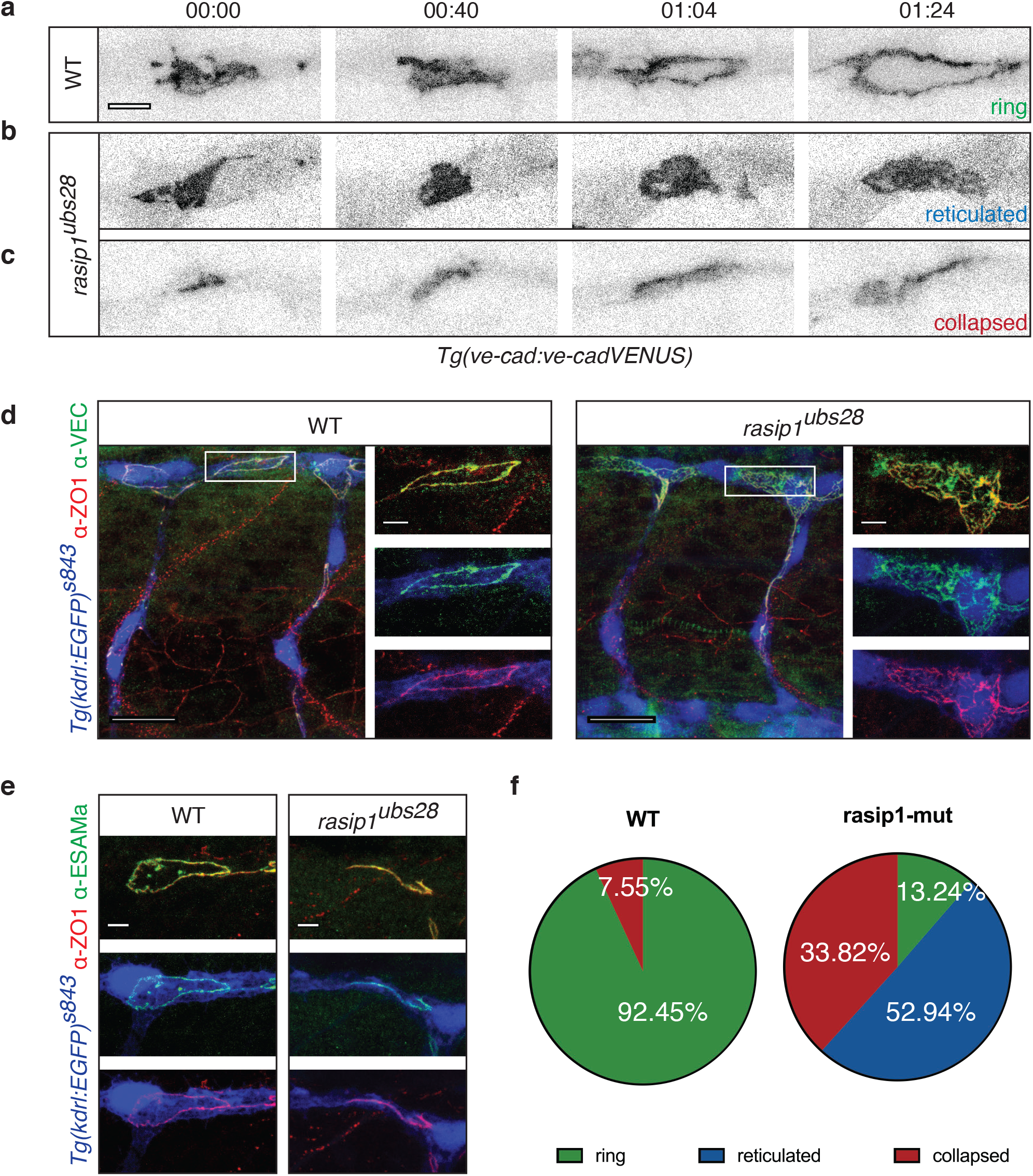
Requirement of Rasip 1 for dynamic re-localization of junctional proteins and junctional ring formation during anastomosis. **(a-c)** Still pictures of time-lapse movies (s-movies 9-11) showing normal junctional patch to ring transformation in wild-type (a) and abberant ring formation in *rasip1*^*ubs28*^ mutants (b, c). Transgenic embryos expressing a VE-cadherin-Venus fusion protein were imaged, starting at 30 hpf. Scale bar, 5 μm. **(d)** Immunofluorescence analysis of Zo-1 and VE-cadherin in *Tg(kdrl:EGFP)*^*s843*^ at 32 hpf. *rasip1*^*ubs28*^ mutants show reticulated junctions between two cells in the DLAV; wild-type embryo forms a cleared, ring-shaped junction. Scale bars, 20 μm (overview) and 5 μm (inset). **(e)** Immunofluorescence analysis of Zo-1 and ESAMa in *Tg(kdrl:EGFP)*^*s843*^ at 32 hpf showing a collapsed junctional ring in *rasip 1*^*ubs28*^ mutants. Scale bars, 5 μm. **(f)** Quantification of observed junctional phenotypes at 32 hpf. *rasip1*^*ubs28*^ mutants show significant number of reticulated junctions and collapsed anastomotic rings compared to wild-type (WT *n*=6 embryos, 53 analyzed rings, mut *n*=8, 68, Chi-Square: p<0.0001.

The described defects were confirmed by immunofluorescent analysis of the junctional proteins VE-cad, ZO-1 (tight junction protein 1, Tjp1) and Esama (endothelial-selective adhesion molecule a) (Figure 3d-e). In *rasip1* mutants VE-cad, Zo-1 as well as Esama colocalized and formed reticulated junctions within the apical compartment. Together, these observations indicate that Rasip1 plays a crucial role in the dynamic re-localization of junctional components during *de novo* junction and lumen formation.

### *rasip1* mutants display transient intracellular luminal pockets

As shown above *rasip1* mutant embryos display reduced vessel diameter and luminal defects (Figure 1e-g). These luminal defects affect the onset of blood flow in the ISVs (Figure 4a, yellow arrowhead). We observed that initially unlumenized ISVs remained unlumenized at least through day 4 of development (96 hpf) (Figure 4b). Moreover, in some instances we observed that initially lumenized blood carrying ISVs collapsed in subsequent stages (up to 96 hpf), indicating a role for Rasip1 in lumen maintenance (Figure 4b,c). Thus, although luminal defects can be attributed to the inability of endothelial cells to rearrange into a multicellular configuration (Figures 2 and 3), the above observations suggest an additional defect in the formation or maintenance of a continuous luminal compartment. This notion is supported by time-lapse analysis of ISV and DLAV formation during lumen formation (Figure 5a). The timing of lumen formation in the ISV is variable but usually starts between 30 and 32 hpf. In wild-type embryos we observed that upon initiation, continuous lumens were formed within 30 minutes (Figure 5a). In *rasip1* mutants, lumen formation was delayed and discontinuous. Instead, we often observed luminal pockets in the dorsal aspects of the ISV (Figure 5a, yellow arrowheads, S-Figure 5d). We surmised that luminal pockets could arise in three different ways (Figure 5b) – either (1) by a collapse of a previously patent tube, (2) by a local cord hollowing event, which failed to interconnect with other luminal pockets or (3) by the formation of large intracellular vacuolar structures, which failed to fuse with luminal membrane.

**Figure 4:**
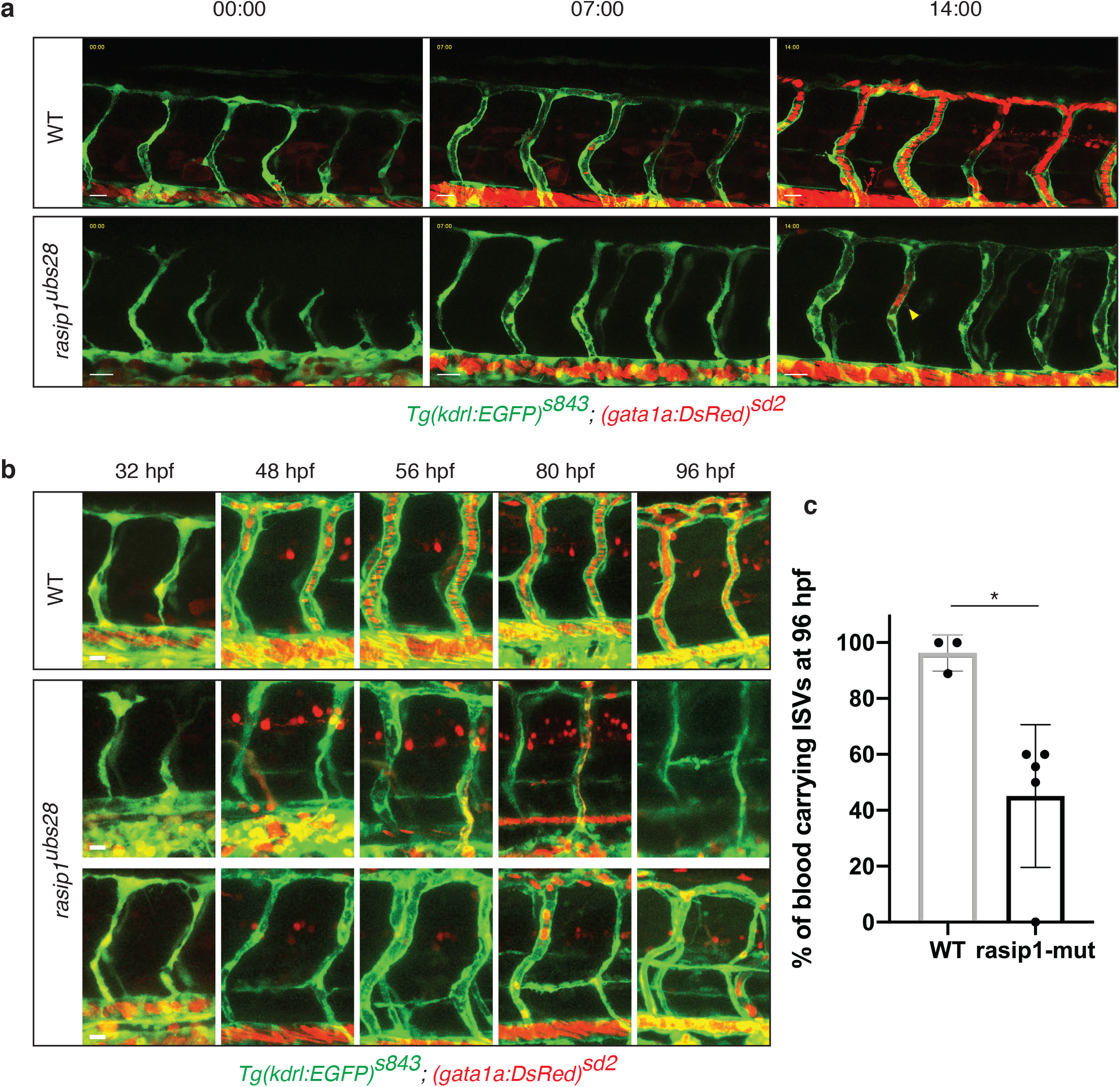
Protracted delays in lumen formation in *rasip1* mutants. **(a, b)** Live images of *Tg(kdrl:EGFP)*^*s843*^; *(gata1a:DsRed)*^*sd2*^ embryos. **(a)** Still images of a time-lapse movies (s-movies 14, 15) starting at 30 hpf. **(b)** Tracking of individual unlumenized ISV during embryonic development (32 to 96 hpf). Scale bars, 20 μm. **(c)** Percentage of blood carrying ISVs at 96 hpf (WT *n*=3 embryos, 28 analyzed ISVs, mut *n*=5, 46). Analyzed by unpaired two-tailed Mann-Whitney test and error bars indicate standard deviation; significance (*p < 0.1).

**Figure 5:**
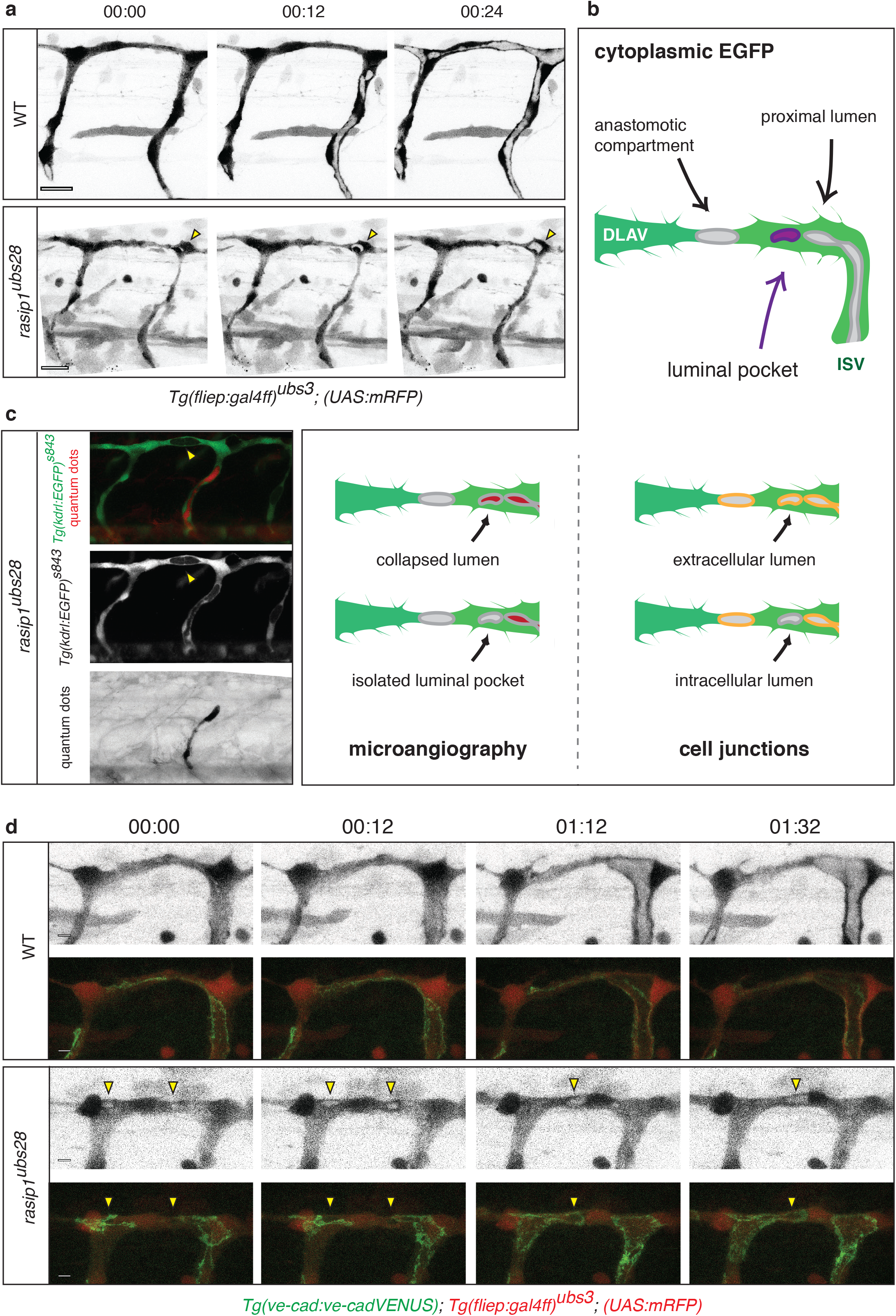
Analysis of ectopic luminal pockets during DLAV formation in *rasip1* mutants. **(a)** Still pictures of time-lapse movies (s-mov 16, 17) showing for the emergence of ectopic luminal pockets (yellow arrowheads) in *rasip1* mutants. **(b)** Schematic representation of possible cellular localization of ectopic lumens. To differentiate between these possibilities two types of experiments: microangiography and colocalization of luminal pockets with junctional marker (d). **(c)** Visualization of ectopic lumens and patent lumens in a *rasip1*^*ubs28*^ embryo (36 hpf). Ectopic luminal pockets are indirectly visualized by the absence of cytoplasmic EGFP (yellow arrowhead) (*Tg(kdrl:EGFP)*^*s843*^). The patent lumen is marked by microangiography using quantum dots in red (third panel in black). Ectopic lumens are not part of the patent vasculature. **(d)** Still pictures of time-lapse movie (s-mov 18-21) during lumen formation in the DLAV from around 32 hpf onward in wild-type (top) and *rasip1*^*ubs28*^ (bottom) embryos. Endothelial cells are labeled with mRFP (greyscale images), junctions are labeled by VE-cad-Venus (merged images). Yellow arrowheads indicate the ectopic luminal pockets in a *rasip1* mutant. Scale bar, 5 μm.

To differentiate between these scenarios, we performed a series of experiments. To test whether luminal pockets may arise by lumen collapse, we performed microangiography using a fluorescent tracer dye in 28 to 30 hpf embryos (Figure 5c). Upon intravascular injection, the entire patent vasculature was labelled by quantum dots. In *rasip1* mutants, however, we observed that although the base of the ISV was positive for quantum dots, local luminal pockets in the DLAV were negative (Figure 5c, yellow arrowheads). This strongly argues that luminal pockets in *rasip1* mutants arise locally and do not represent luminal remainders formed by lumen collapse.

We next wanted to check whether luminal pockets represented intra- or extracellular compartments. Extracellular luminal compartments arise between cells in a process called cord hollowing, whereas intracellular lumens are thought to form by vacuolation (Davis et al., 2011). During anastomosis in zebrafish embryos, cord hollowing generates luminal pockets form as transient structures at the interface between contacting tip cells (Blum et al., 2008). This interface is formed by a ring-shaped junction, which surrounds an apical compartment of both tip cells (Herwig et al., 2011). Thus, these extracellular pockets are demarcated by junctional rings, while intracellular pockets should be outside of these rings. To test these possibilities, we examined lumen formation in *rasip1* mutants expressing VE-cad-Venus and found that they were located outside the junctional ring (Figure 5d (00:12)), appearing as vesicular structures within the endothelial cytoplasm. Lateron (Figure 5d (01:32)) these intracellular lumens were incorporated into the area covered by the junctional ring, therefore representing transient structures. Taken together, these findings show that loss of Rasip1 function leads to a transient accumulation of intracellular vesicles, which later on merge into the anastomotic compartment suggesting that Rasip1 may be critical for normal vesicle transport or vesicle fusion during cord hollowing process, which occurs during anastomosis.

### Rasip1 localizes to apical membranes and endothelial cell junctions

Our mutant analyses indicate a requirement for Rasip1 in junction formation and remodeling, as well as in lumen formation and maintenance. To gain a better understanding of how Rasip1 may be involved in these processes, we generated an antibody against zebrafish Rasip1 to discern the subcellular localization of the protein during angiogenesis. Immunofluorescent analysis confirmed endothelial expression of Rasip1 in zebrafish embryos (S-Figure 4). Notably, Rasip1 protein levels appeared dynamically regulated. During vasculogenesis, until the emergence of intersegmental sprouts, Rasip1 was detected at high levels in the dorsal aorta (S-Figure 4a,b, yellow arrowheads). In contrast, Rasip1 protein was highly expressed in sprouting ISVs while it became downregulated in the dorsal aorta (S-Figure 4c, yellow bars), supporting the notion that Rasip1 primarily functions during blood vessel morphogenesis rather than during vessel maintenance. High-resolution imaging revealed specific subcellular localization during blood vessel formation. In the context of anastomosis, three different phases could be discerned. First, during contact formation (30 hpf), we found that Rasip1 is absent from newly formed contacts (Figure 6a, yellow arrowheads). However, Rasip1 was observed colocalizing with large junctional patches prior to discernable formation of apical compartments. During later stages of anastomosis, when the anastomotic ring had formed, Rasip1 was restricted to the apical compartment within the junctional ring with no detectable Rasip1 at the junction (Figure 6b, yellow arrowheads). However, shortly later - during the establishment of the DLAV (36 hpf) and later after full development of angiogenic vessels (48 hpf) -, we found that Rasip1 also localized to endothelial cell junctions (Figure 6c, white arrowheads). Taken together, these studies show that Rasip1 dynamically distributes during different phases of blood vessel formation. In particular, the dynamic subcellular distribution to apical membrane compartments and endothelial cell junctions suggests a sequential requirement for Rasip1 during apical compartment formation and junctional remodeling, respectively.

**Figure 6:**
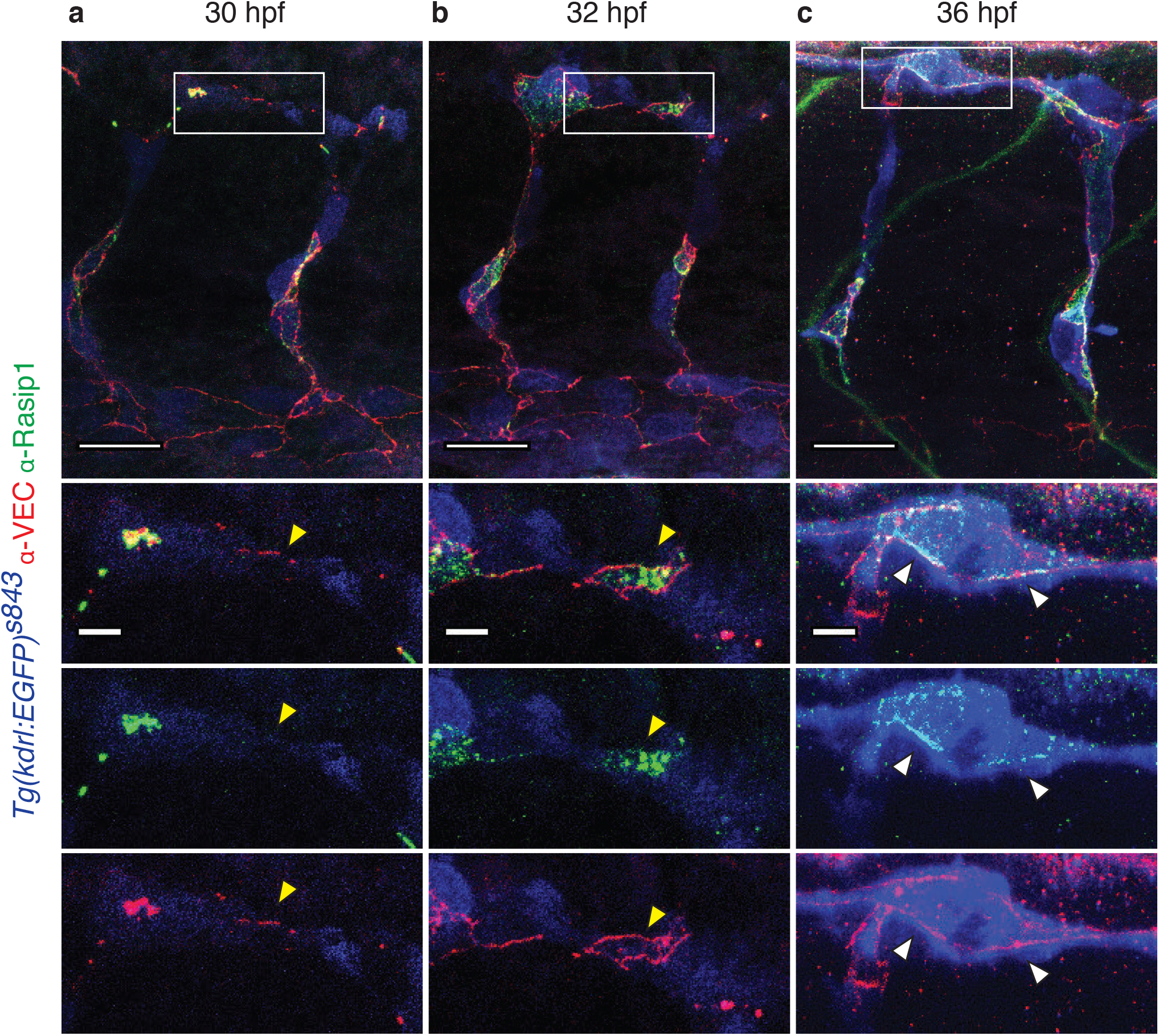
Apical to junctional re-localization of Rasip1 during blood vessel fusion. **(a-c)** Immunofluorescent labeling of Rasip1 and VE-cadherin during different stages of DLAV formation (30 to 36 hpf). At 32hpf, Rasip1 is restricted to the apical surface of the anastomotic ring (yellow arrowheads). At 36hpf, Rasip1 localizes to endothelial junctions (VE-cadherin, white arrowheads). Scale bars, 20 μm (overview) and 5 μm (inset).

Because of the early localization of Rasip1 to the apical membrane, we wanted to test whether loss of Rasip1 function may affect apical polarization during blood vessel formation. To this end, we generated a transgenic reporter (*Tg(EGFP-podxl)*^*ubs29*^), which labels the apical membrane compartment (S-Figure 5a). At 48 hpf, we observed normal luminal localization of EGFP-Podxl in *rasip1* mutants (S-Figure 5b). In ISVs that displayed luminal defects, we observed slightly irregular distribution of EGFP-Podxl in affected areas (enlarged insets in S-Figure 5b). These observations suggest that in spite of its apical localization, Rasip1 is not required for apical polarization in endothelial cells.

### Overlapping requirement of Rasip1 and Radil-b in blood vessel formation

Rasip1 protein has been shown to be an effector protein of the small GTPase Rap1 (Gingras et al., 2016). Protein binding studies have shown that Rasip1 can form multimeric complexes consisting of Rap1, Rasip1, Radil (Ras-associating-dilute-domain) and the GTPase-activating protein Arhgap29 (Post et al., 2013). Furthermore, it has been shown that the core of these complexes can be formed by a Rasip1 homodimer or a Rasip1/Radil heterodimer (de Kreuk et al., 2016), indicating partly overlapping functions of these proteins in endothelial cells. Rasip1 and Radil are closely related proteins (S-Figure 6a) sharing several protein interaction domains such as a Ras association (RA) domain conferring binding to Rap1, and a forkhead association (FHA) domain binding the transmembrane receptor Heg1. An additional PDZ domain is unique to Radil and is thought to interact with the GTPase-activating protein Arhgap29 (Post et al., 2013). Radil function in endothelial cells has, so far, only been addressed in cell culture experiments, which established its above-mentioned protein interactions and indicated a role of Radil in endothelial barrier maintenance and the regulation of endothelial cell adhesion (de Kreuk et al., 2016; Pannekoek et al., 2014). Thus, we wanted to determine the role of Radil during blood vessel morphogenesis *in vivo* and compare its requirement to that of Rasip1. The zebrafish genome contains three *radil* paralogues, *radil-a,-b* and -*c*, respectively (S-Figure 6a). Whole-mount in situ analysis revealed that of the three paralogues, only *radil-b* was expressed specifically in endothelial cells (data not shown). Therefore, we analyzed blood vessel formation in *radilb*^*sa20161*^ mutants, which carry a nonsense mutation (Tyr129 → STOP) near the N-terminus of the protein (S-Figure 6b).

*radil-b* mutants were homozygous viable and could be raised to fertility. Nevertheless, they exhibited several vascular defects similar to *rasip1* mutants, including cerebral hemorrhages (S-Figure 6c), isolated luminal pockets (S-Figure 6d), reduction of blood flow (S-Figure 6e) and retarded sprouting of ISVs (data not shown), supporting the notion that both proteins are involved in the same molecular pathways. In contrast, *radil-b* mutants did not phenocopy the junctional re-localization defect observed in *rasip1* mutants during anastomosis (Figure 7a), suggesting that the proteins may also have unique functions. *radil-b* mutants generally exhibited milder defects (S-Figure 6f, g), in particular with respect to cellular architecture compared to single *rasip1* mutants and *rasip1;radil-b* double mutants (Figure 7b). Furthermore, *rasip1;radil-b* double mutants showed stronger sprouting and lumen formation defects than either single mutant (Figure 7, S-Figure 6h), suggesting that while both proteins are required in this process, they likely act in a partially redundant manner.

**Figure 7:**
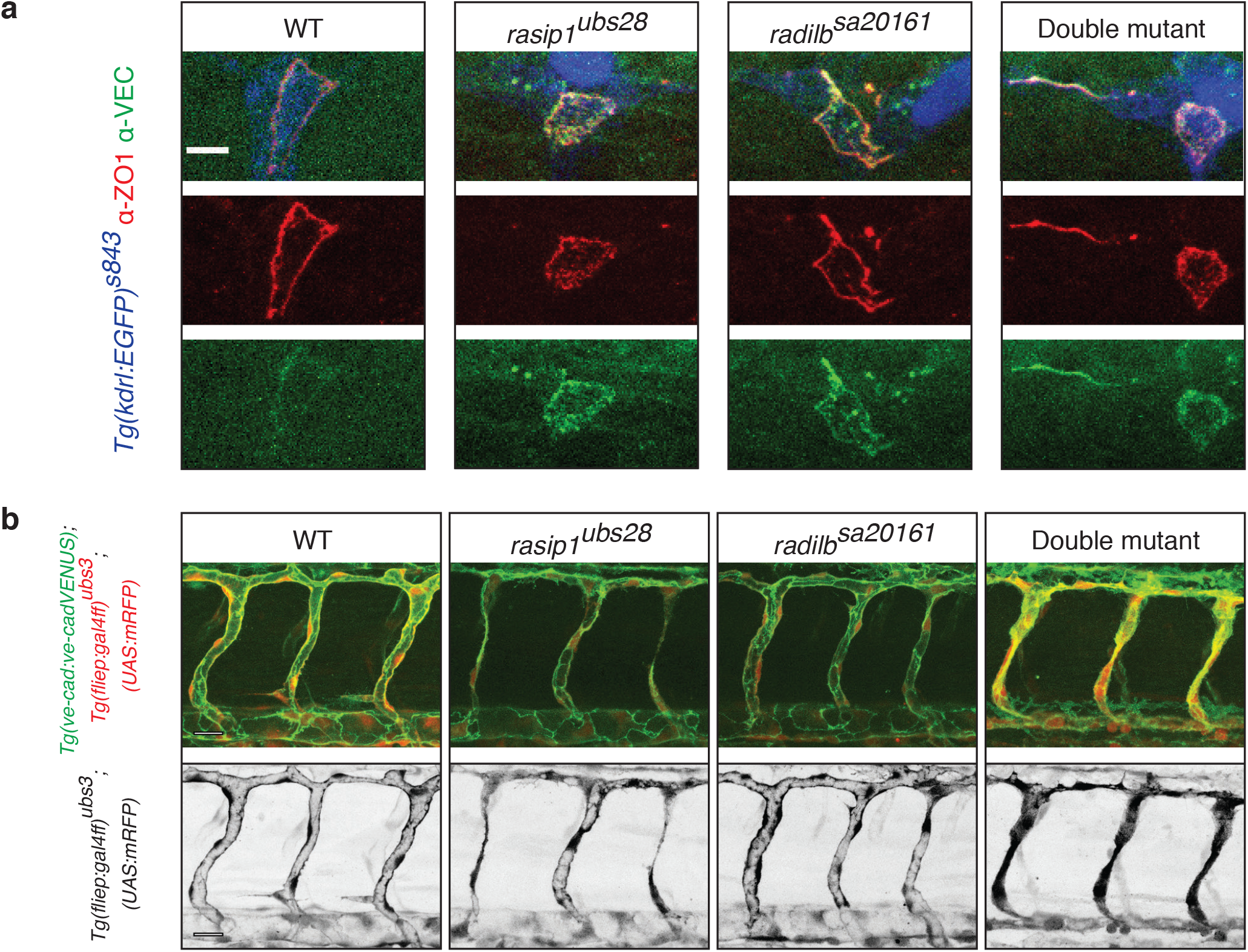
Phenotypic comparison *of radil-b* single and of *rasip1/radil-b* double mutants suggests partially overlapping functions during vascular morphogenesis. **(a)** Immunofluorescence analysis of Zo-1 and VE-cadherin distribution in *Tg(kdrl:EGFP)*^*s843*^ at 32 hpf. **(b)** Live images at 48 hpf using *Tg(ve-cad:ve-cadVENUS); Tg(fliep:gal4ff)*^*ubs3*^; *(UAS:mRFP*) reporter lines. Scale bars: 20 µm

### Knock-down of *ccm1* and *heg1* phenocopies aspects of the *rasip1* mutant

The Rasip1/Radil/Rap1 complex can bind via the FHA domain to the orphan transmembrane receptor Heg1 (Gingras et al., 2016). This interaction has been shown to tether Rasip1 to endothelial cell junctions (Post et al., 2013). To address the relevance of this interaction during ISV morphogenesis, we analyzed *heg1* morphants in order to test whether loss of Heg1 function showed any *rasip1*-like vascular phenotypes. We also examined the phenotypes of *ccm1* morphants. Ccm1 binds to Heg1 independently of Rasip1 (Gingras et al., 2012), and thus might indirectly influence Rasip1 function. *ccm1* and *heg1* morphants exhibited blood vessel and heart dilations as previously published (S-Figure 7a) (Hogan et al., 2008; Kleaveland et al., 2009; Stainier et al., 1996). We consistently observed hemorrhages, which were less pronounced than in *rasip1* mutants (Figure 8a and S-Figure 7b). However, similar to *rasip1* mutants, ISV sprouting and development appeared to be delayed (Figure 8b). To assess the role of Heg1 and Ccm1 in junctional rearrangements, we analyzed endothelial cell junctions in transgenic VE-cad-Venus and Pecam1-EGFP reporter background (S-Figure 7e). Consistent with earlier reports (Hogan et al., 2008; Kleaveland et al., 2009), knock-down of *ccm1* or *heg1* did not cause obvious junctional defects in the dorsal aorta. However, we observed detachment of junctions and impaired cell rearrangements in sprouting ISVs (S-Figure 7c, d). In addition, it appeared that Pecam-EGFP as well as VE-cad-Venus were not entirely cleared from apical compartments during the formation of the DLAV.

**Figure 8:**
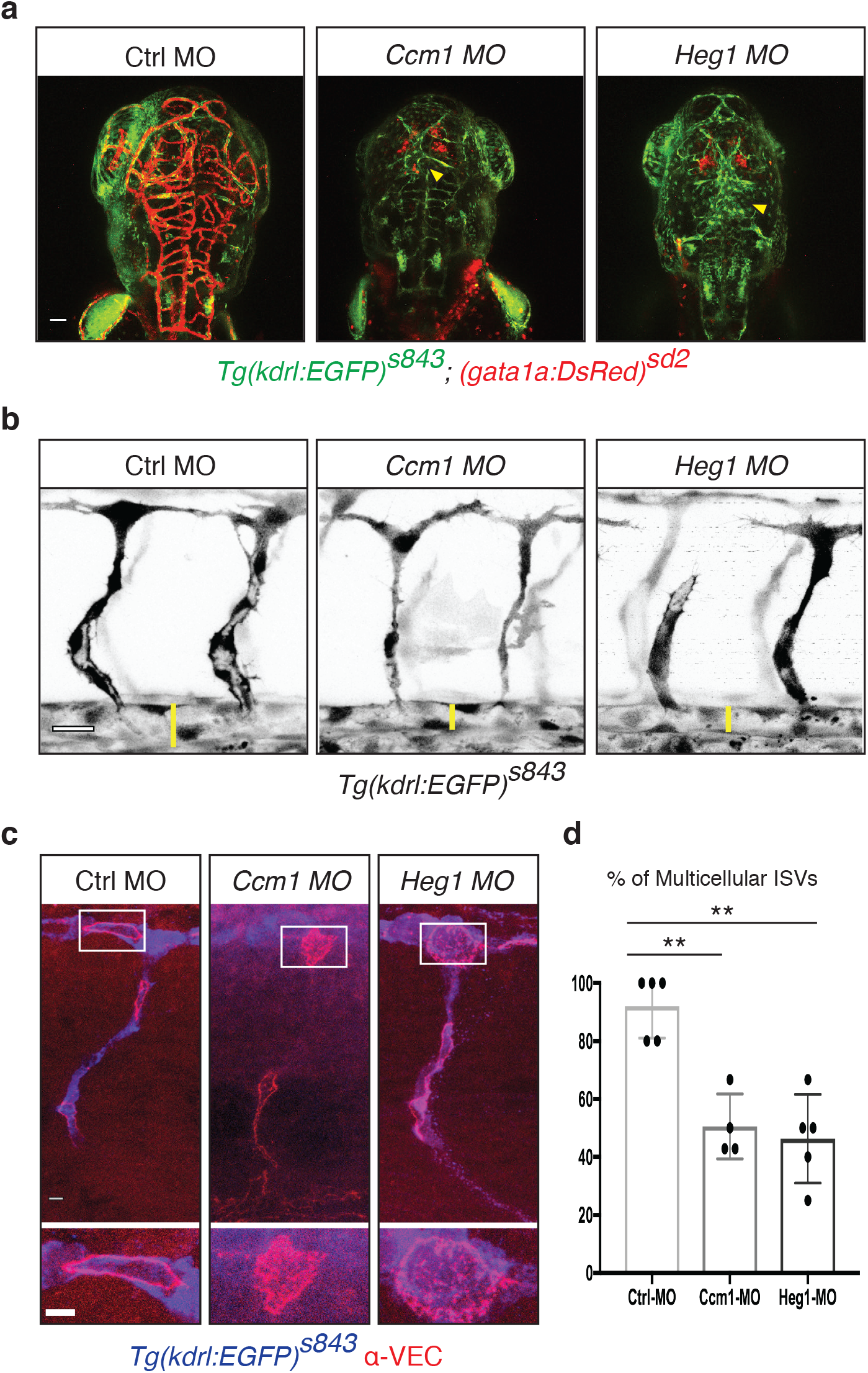
Loss of Ccm1 and Heg1 phenocopies aspects of *rasip1* mutants. **(a)** Live images of Tg(kdrl:EGFP)^s843^; (gata1:Dsred)^sd2^ at 72 hpf. *ccm1* and *heg1* morphants display mesenphalic hemorrhages while cranial circulation appears completely disrupted. In addition, the MsV (Mesencephalic veins) and DLV (dorsal longitudinal vein) (yellow arrowheads) are malformed. Scale bar, 50 μm. **(b)** Live images of control, *ccm1* and *heg1* morphants at 32 hpf. *ccm1* and *heg1* morphants show reduced DA diameters and defective ISV formation. Scale bar, 20 μm. **(c)** Immunofluorescence analysis of control, *Ccm1* and *Heg1* morphants at 32 hpf. Transgenic *Tg(kdrl:EGFP)*^*s843*^ embryos were stained for VE-cadherin. Scale bar, 5 μm. **(d)** Quantification of multicellular ISVs at 48 hpf (Control MO injected embryos *n*=5, 24 analyzed ISVs; *Ccm1* MO *n*=4, 26; *Heg1* MO *n*=5, 23). Analyzed by unpaired two-tailed Mann-Whitney test and error bars indicate standard deviation; significance (**p < 0.01).

To verify these observations, we performed immunofluorescent analyses to determine whether endogenous VE-cadherin protein was cleared from the apical membrane compartment in *heg1* and *ccm1* morphants. In both conditions, VE-cadherin accumulated in the apical compartment embedded within the junctional rings – similar to what we observed of *rasip1* mutants (Figure 8c). *In vivo* imaging of the junctions in *ccm1* or *heg1* knock-down embryos showed a failure of multicellular vessel formation (Figure 8d). Taken together, these findings show that *ccm1* and *heg1* loss-of-function phenocopy defined aspects of *rasip1* mutants during critical aspects of blood vessel formation during sprouting angiogenesis and anastomosis.

## Discussion

Small GTPases of the Rho family play a key role in the regulation of cellular activities during blood vessel formation. For example, they serve as molecular switches to control cytoskeletal dynamics, cell adhesion and junction assembly during angiogenic sprouting and lumen formation (Barlow and Cleaver, 2019). Rasip1 has been described as an effector protein of small GTPase signaling during blood vessel formation and maintenance (Koo et al., 2016; Wilson et al., 2013; Xu et al., 2011). Rasip1 protein contains multiple protein binding domains and has been shown to directly interact with its paralogue Radil, the small GTPase Rap1 and the transmembrane protein Heg1 (de Kreuk et al., 2016; Gingras et al., 2016; Wilson et al., 2013). By further association with proteins such as Arhgap29 and Ccm1, these proteins control cortical actomyosin tension and endothelial junction formation and dynamics (Post et al., 2015) and reviewed by (Lampugnani et al., 2017; Wilson and Ye, 2014). Analyses of mutant mouse embryos have shown that Rasip1 is required for proper lumen formation and maintenance in blood and lymphatic vessels (Koo et al., 2016; Liu et al., 2018; Wilson et al., 2013). During vasculogenesis, Rasip1 is required for the establishment of the nascent apical compartment and subsequently for lumen expansion, presumably mediated by regulating Cdc42 and RhoA, respectively (Barry et al., 2016).

### Multiple vascular defects in zebrafish *rasip1* mutants

In order to gain a better understanding of how morphogenetic cell behaviors are controlled by Rasip1, we generated loss-of-function mutants in the zebrafish *rasip1* gene and analyzed these mutants focusing on cellular and junctional dynamics during angiogenic sprouting and anastomosis. Overall, our findings are in agreement with previously published vascular defects in mouse development, but also provide novel insights into the regulation of junctional dynamics during angiogenic sprouting and lumen formation.

*rasip1* mutants display numerous vascular defects, including cranial hemorrhage, reduced blood circulation and reduced diameter of the dorsal aorta, consistent with previously published *rasip1* knockdown experiments in zebrafish (Wilson et al., 2013). Furthermore, we observe delayed angiogenic sprouting, as well as abnormalities in lumen formation as well as impaired cell rearrangements, junctional dynamics and stability. Despite this wide range of defects, the mutant phenotypes are consistent with defects in the control of F-actin and junctional dynamics. The phenotypes of *rasip1* mutants during vascular development are, however, quite distinct from those seen in *ve-cad (cdh5)* mutants (Sauteur et al., 2014) indicating that Rasip1 is regulating endothelial activities beyond cell junctions.

### Dynamic regulation of Rasip1 expression and subcellular localization

Endothelial-specific expression of Rasip1 has been reported in several vertebrate species including mouse, *Xenopus* and zebrafish, and Rasip1 appears to be expressed in the entire embryonic vasculature (Wilson et al., 2013; Xu et al., 2009). In contrast to its broad endothelial transcription, the distribution of Rasip1 protein appears to highly regulated, with respect to the overall level and its subcellular localization. During vasculogenesis, Rasip1 is readily detected at high levels in the dorsal aorta, whereas at later stages expression is somewhat reduced. This downregulation of Rasip1 in the dorsal aorta coincides with the commencement of blood flow, suggesting that Rasip1 protein levels and localization may be controlled by shear stress. The downregulation of Rasip1 protein in more mature vessels suggests an essential role during blood vessel morphogenesis. In agreement with this, Rasip1 does not appear to be required in established blood or lymphatic vessels (Koo et al., 2016; Liu et al., 2018).

During ISV sprouting we observe a shift of Rasip1 from the apical membrane compartment during the early stages to higher levels at cell junctions during the later stages of vascular tube formation. The early apical localization of Rasip1 is in agreement with a previously proposed role in early apical-basal polarization, likely upstream of Cdc42 (Barry et al., 2016). It will be important to determine how this differential localization is accomplished or to what extend it reflects different functions of Rasip1 during tubulogenesis. In *Drosophila*, it has been shown that apical localization of canoe (the ortholog of vertebrate *afadin* and a homologue of rasip1) is dependent on Rap1 (Bonello et al., 2018). Therefore, Rap1 is a putative candidate to localize Rasip1 to apical membrane during vertebrate angiogenesis.

### A role for Rasip1 in angiogenic sprouting and blood vessel assembly

Sprouting angiogenesis is accomplished by concerted endothelial cell dynamics including cell migration, rearrangement, elongation and proliferation, all of which are affected by the loss Rasip1 function. *rasip1* mutant ISVs contain fewer cells than wild-type ISV, which reflects a reduced rate of cell proliferation during sprouting.

Furthermore, we observe a frequent failure of *rasip1* mutant ISVs to form multicellular tubes. Formation of multicellular ISVs does not depend of the number of cells present in the sprout (Angulo-Urarte et al., 2018) but rather relies on cell rearrangements driven by junctional remodeling (Sauteur et al., 2014). For example, loss of VE-cad prevents junction elongation and renders endothelial cells unable to move over each other and effectively pair to form a multicellular tube (Paatero et al., 2018; Sauteur et al., 2014). In *rasip1* mutants, however, the defect in cell pairing is caused by junctional detachments. When we imaged VE-cad dynamics during ISV sprouting in wild-type embryos, we found that, in many cases, two stalk cells maintain contact to the dorsal aorta. In *rasip1* mutant sprouts, we observed that one of these cells detached from the dorsal aorta and “retracted” to the dorsal part of the ISV, leaving the remaining cell unpaired. Stalk cell detachment appears to occur at tri-cellular junctions indicating that these junctions may be important as an anchor point to resist mechanical forces occuring during cell rearrangements. A study performed in MDCK cells has shown that Afadin accumulates at tri-cellular junctions in response to tension (Choi et al., 2016). This finding supports the interesting notion that Rasip1 may play a specific role in reinforcing tri-cellular junctions during sprouting angiogenesis.

### Rasip1 is required for lumen patency

Lumen formation in *rasip1* mutants is delayed and ISVs as well as the dorsal aorta show reduced vessel diameters during early embryonic development. At 48 hpf, we observed that about 50% of the ISVs were not patent and this defect was maintained at least through day 4 of development. Defects in luminal patency may have multiple causes. First, as described above, junctional detachment can prevent endothelial cell pairing and thus multicellular tube formation. Second, our examinations of the apical membrane using a Podxl-EGFP and Cherry-CAAX transgenic markers revealed an irregular shape of the apical membrane, suggesting luminal collapse. This observation agrees with earlier findings that Rasip1 regulates cortical actin tension during lumen opening of the dorsal aorta in mouse (Barry et al., 2016).

During the formation of the DLAV formation, we also observed luminal pockets, which did not label by microangiography and appeared outside of cell junctions, indicating that these lumens are intracellular and consist of large vesicles or vacuolar structures. Rasip1 has been shown to associate with early Rab5-positive and recycling Rab8-positive endosomes (Barry et al., 2016) and Rab8 has been implicated in the transport of Podocalyxin to the apical membrane in a Cdc42 dependent manner (Bryant et al., 2010). Further analyses will have to be undertaken to determine whether these intracellular lumens are endosomal compartments and whether Rasip1 plays a role in targeting recycling endosomes to the apical compartment.

### Rasip1 during anastomosis

Vascular anastomosis is the process by which blood vessel connect and form a network. Formation of the DLAV in the zebrafish is initiated by the interaction of two neighboring tip cells which establish contact and form a localized *de novo* lumen at their interface (Herwig et al., 2011) reviewed by (Betz et al., 2016). Formation of this luminal pocket follows a relatively stereotyped sequence: upon initial interfilopodial contact a junctional spot is formed, which is transformed into a ring surrounding apical membrane. This spot to ring transformation entails the formation and expansion of a stable junctional ring and the removal of junctional proteins from the center to permit formation of an apical membrane compartment. Loss of Rasip1 prevents apical clearance leading to ectopic junctions within newly formed apical compartments. A similar phenotype has been observed in *Rasip1* mouse mutants during lumen formation of the dorsal aorta and it was shown that this requirement is upstream of Cdc42 (Barry et al., 2016). Thus, the molecular mechanisms driving apical clearance during vasculogenesis and anastomosis appear to be rather conserved.

*In vitro* experiments have shown that Rasip1 localization at cell junctions requires its interaction with the orphan receptor Heg1 (de Kreuk et al., 2016) and it has been suggested that Rasip1 acts in concert with Heg1, Rap1 and Ccm1 and other proteins in junction stabilization (reviewed by (Lampugnani et al., 2017)). To test whether Rasip1, Ccm1 and Heg1 may interact during apical clearance, we performed knockdown experiments. Knockdown of *ccm1* as well as *heg1* in zebrafish phenocopy the apical clearance defects seen in *rasip1* mutants. Taken together, these data suggest that some of the pathways which are involved in junction stabilization are also required for or result in apical clearance during *de novo* lumen formation.

Rasip1, Radil and Arghap29 have been shown to form a complex and are thought to regulate RhoA. Our analysis of *radil-b* and *rasip1/radil-b* double mutants has shown that both proteins have similar functions during angiogenic sprouting and lumen formation and maintenance. However, Radil-b appears to be dispensable for apical junctional re-localization during anastomosis. In agreement with this interpretation, studies in the mouse dorsal aorta have shown that clearance of apical junction requires Cdc42 and is independent of RhoA (Barry et al., 2016).

In mouse dorsal aortae, loss of Rasip1 leads to an overactivation of Rock and an increase of cortical actomyosin tension in the apical compartment (Barry et al., 2016). As a consequence, the luminal surface of the endothelium cannot expand and the lumen is constricted. In relation to this, we observe a collapse of the junctional ring during anastomosis while the junctions are still maintained and elongate along the vascular axis, resulting in a narrower apical compartment within the ring, eventually leading to a close alignment of the junctions along the extending axis. We speculate that this collapse of the junctional ring may be caused by an imbalance of the cortical actin between apical and basolateral compartment, caused by an overactivation of Rock at the apical side. Further studies on local actomyosin regulation will be required to better understand the formation of a luminal surface during vasculogenesis and vascular anastomosis.

## Materials and Methods

### Zebrafish Strains and Morpholinos

Zebrafish were maintained according to FELASA guidelines (Aleström et al., 2019). All experiments were performed in accordance with federal guidelines and were approved by the Kantonales Veterinäramt of Kanton Basel-Stadt. Zebrafish lines used were *Tg(gata1a:DsRed)*^sd2^ (Traver et al., 2003), *Tg(kdrl:EGFP)*^*s843*^ (Jin, 2005), *Tg(kdrl:EGFPnls)*^*ubs1*^ (Blum et al., 2008), *Tg(5xUAS:RFP)* (Asakawa and Kawakami, 2008), *Tg(fli1ep:gal4ff)*^*ubs3*^ (Herwig et al., 2011), *Tg(fli1a:Pecam-EGFP)*^*ncv27*^ (Ando et al., 2016), *Tg(cdh5:cdh5-TFP-TENS-Venus)*^*uq11bh*^ (Lagendijk et al., 2017), *Tg(UAS:EGFPpodxl)*^*ubs29*^ (this study) and *rasip1*^*ubs28*^ (this study) and *radilb*^*sa20161*^ (European Zebrafish Resource Center, Karlsruhe, Germany). Morpholinos (Gene-Tools, Corvallis, OR, USA) used were as follows: *ccm1* 5’-GCTTTATTTCACCTCACCTCATAGG-3’ (Mably, 2006), *heg1* 5’-GTAATCGTACTTGCAGCAGGTGACA-3’ (Mably et al., 2003), standard control 5’-CCTCTTACCTCAGTTACAATTTATA-3’.

### Generation of *Tg(UAS:EGFPpodxl)*^*ubs29*^

The p5E-4xnrUAS promoter, pME-EGFP-podocalyxin (Navis et al., 2013) and p3E-polyA (Kwan et al., 2007) were cloned into a Tol2 vector pDestTol2CG2 carrying cmcl2:GFP to drive expression of GFP in the heart. The final plasmid was co-injected with tol2 mRNA into the *Tg(fli1ep:gal4ff)*^*ubs3*^; *(UAS:mRFP)* embryos. These mosaic embryos were raised to adulthood and outcrossed with the parental fish line to generate stable fish lines. The resulting *Tg(UAS:EGFPpodxl)* embryos were identified on the basis of GFP expression in the heart; proper apical localization of EGFP-Podocalyxin was confirmed using confocal microscopy. Two transgenic lines, ubs29 and ubs30, were isolated, and the ubs29 line showing more homogenous expression levels in endothelium was used in experiments.

### Immunofluorescence

Immunofluorescence was performed as previously described (Herwig et al., 2011). The following antibodies were used: rabbit anti-zf-Cdh5 1:200 (Blum et al., 2008), rabbit anti-Esama 1:200 (Sauteur et al., 2017), mouse anti-human-Zo-1 1:100 (Thermofisher), rabbit anti-Rasip1 1:500 (this paper), chicken anti-GFP 1:200 (Abcam), Alexa 405 goat anti-chicken immunoglobulin Y (IgY H&L) 1:1,000 (Abcam), Alexa 568 goat anti-rabbit immunoglobulin G (IgG) 1:1,000, and Alexa 633 goat anti-mouse IgG 1:1,000 (both from Thermofisher). The anti-zf-Rasip1 antibodies were raised in rabbits against a synthetic peptide (CRTFLWGLDQDELPANQRTRL-COOH) comprising the terminal amino acid residues (aa970-989) of the protein (Yenzym, Antibodies LLC, Brisbane (CA, USA)).

### Live Imaging

Time-lapse imaging was performed as previously described (Paatero et al., 2018). All movies were taken with Leica SP5 or SP8 confocal microscopes using a 40x water immersion objective (NA = 1.1) with a frame size of 1024×512 or 1024×1024 pixels. Routinely, z stacks consisted of 80–100 slices with a step size of 0.8–1 μm. Stacks were taken every 8- or 10-min. High-resolution imaging was performed a Zeiss LSM880 microscope using a 40x water immersion objective (NA = 1.2) using a vertical step size of 0.25 µm.

### Statistics

Unless explicitly stated, all results shown were obtained from at least 3 independent experiments, sample sizes were not predetermined, the experiments were not randomized and investigators were not blinded to allocation during experiments and outcome assessment. Statistical analyses were performed using Prism software (GraphPad) and ordinary unpaired two-tailed Mann-Whitney test.

## Author Contributions

H.-G.B. and M.A. conceived the project; M.L. performed most experiments and prepared the figures. C.B. generated *rasip1* mutant alleles and performed initial experiments. N.S. characterized *radil-b* mutants; I.P. generated the *Tg(UAS:EGFPpodxl)*^*ubs29*^ zebrafish line; J.Y. analyzed *heg1* and *ccm1* morphants; C.W.W. and W.Y. provided the anti-zf-Rasip1 antibody; M.A and H.-G.B. supervised the project. M.L., M.A. and H.-G.B. wrote the manuscript. All authors read and approved the manuscript.

## Acknowledgements

We would like to thank the Biozentrum Imaging Core Facility for ceaseless support, Dr. Li-Kun Phng and Gustavo Aguilar for critically reading the manuscript, Dr. Anne Karine Lagendijk and Dr. Benjamin M. Hogan for providing the transgenic VE-cad-Venus reporter, Dr. David Dylus for help on phylotypic analysis and Kumuthini Kulendra and Mattias Thimm for fish husbandry. M.L. was supported by a fellowship from the Werner-Siemens-Foundation (Zug). This work has been supported by the Kantons Basel-Stadt and Basel-Land and by a grant from the Swiss National Science Foundation to M.A.

