## Supplementary Information for "Control of dynamic cell behaviors during angiogenesis and anastomosis by Rasip 1"

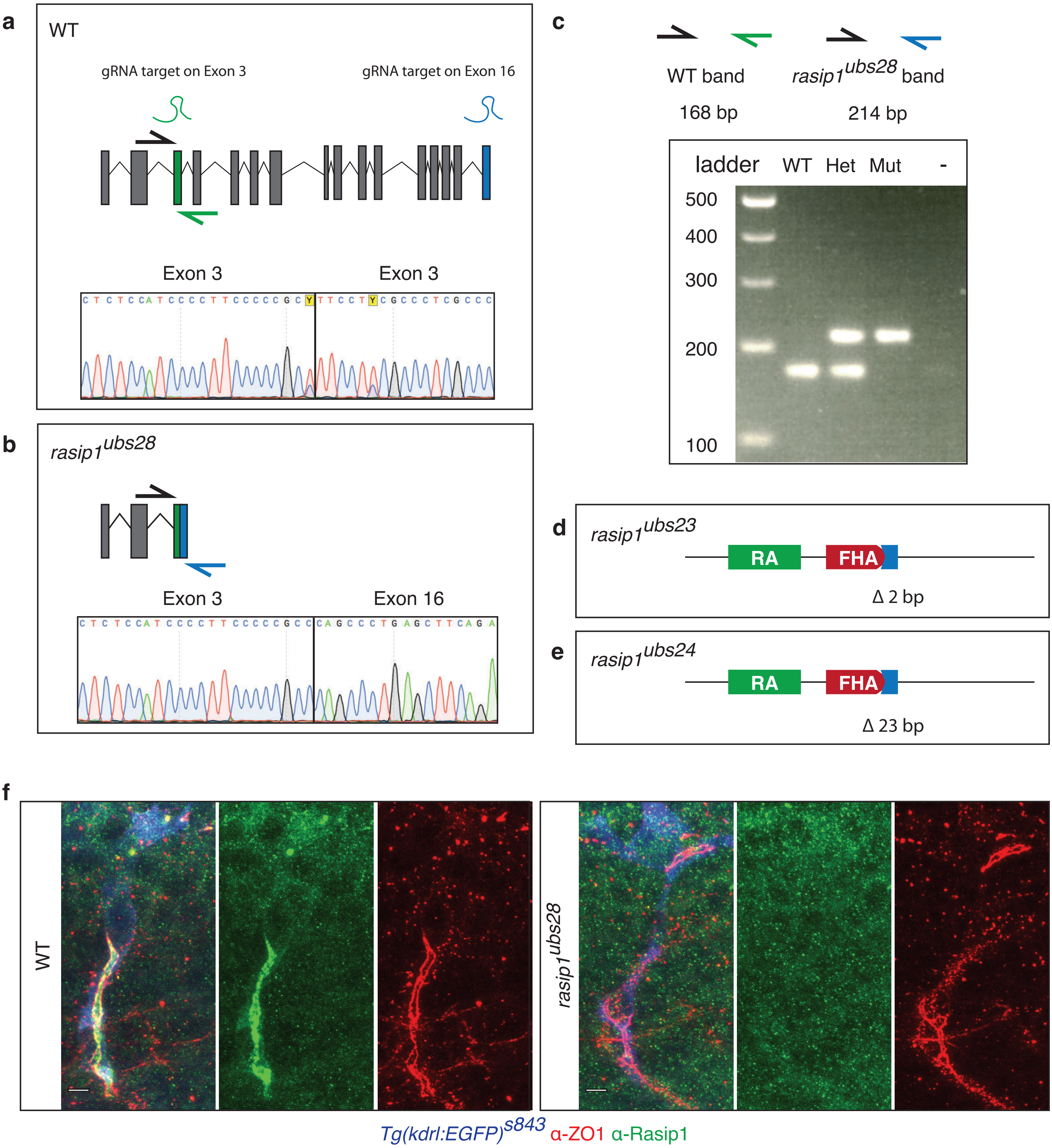


Supplementary Figure 1

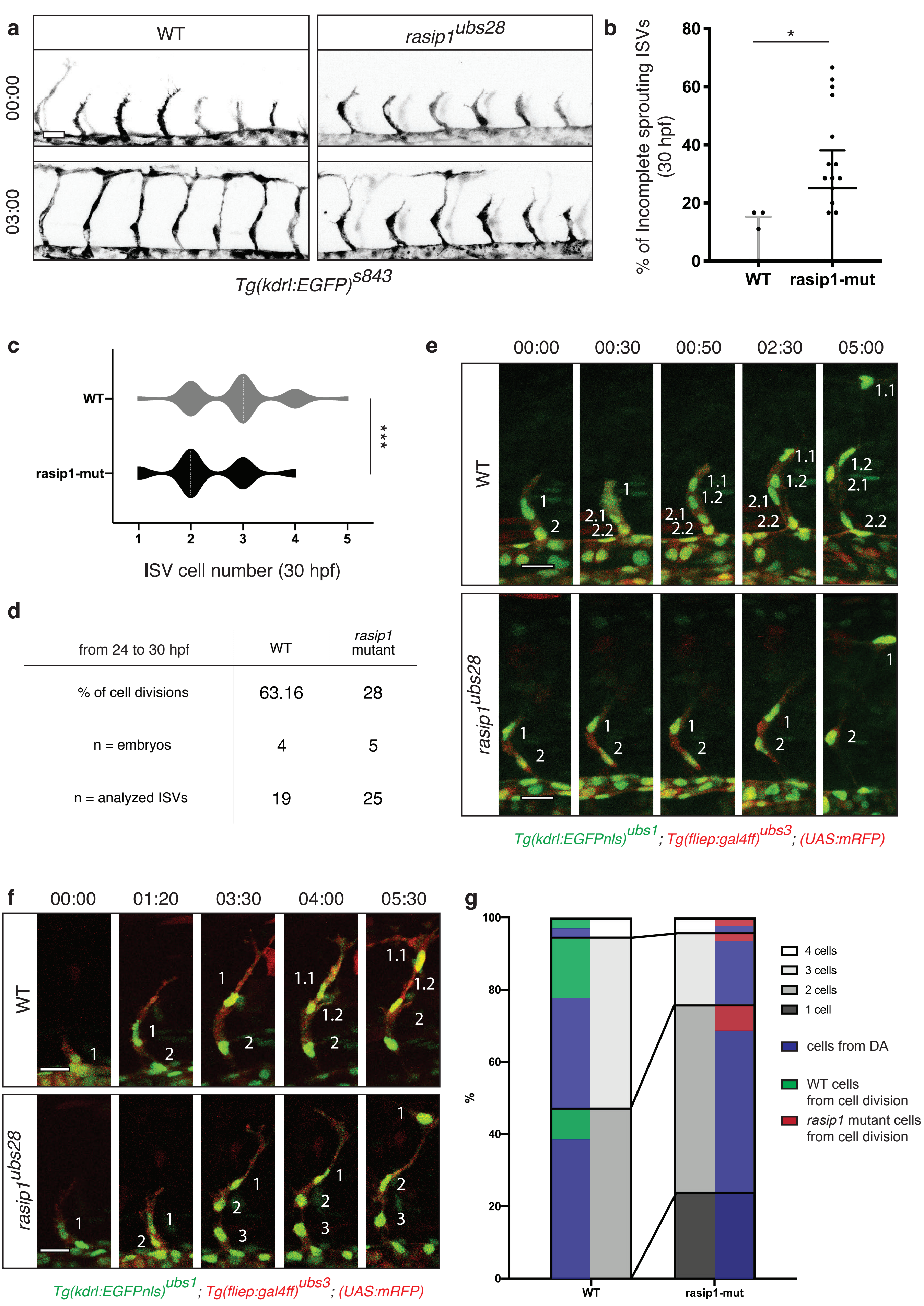


Supplementary Figure 2

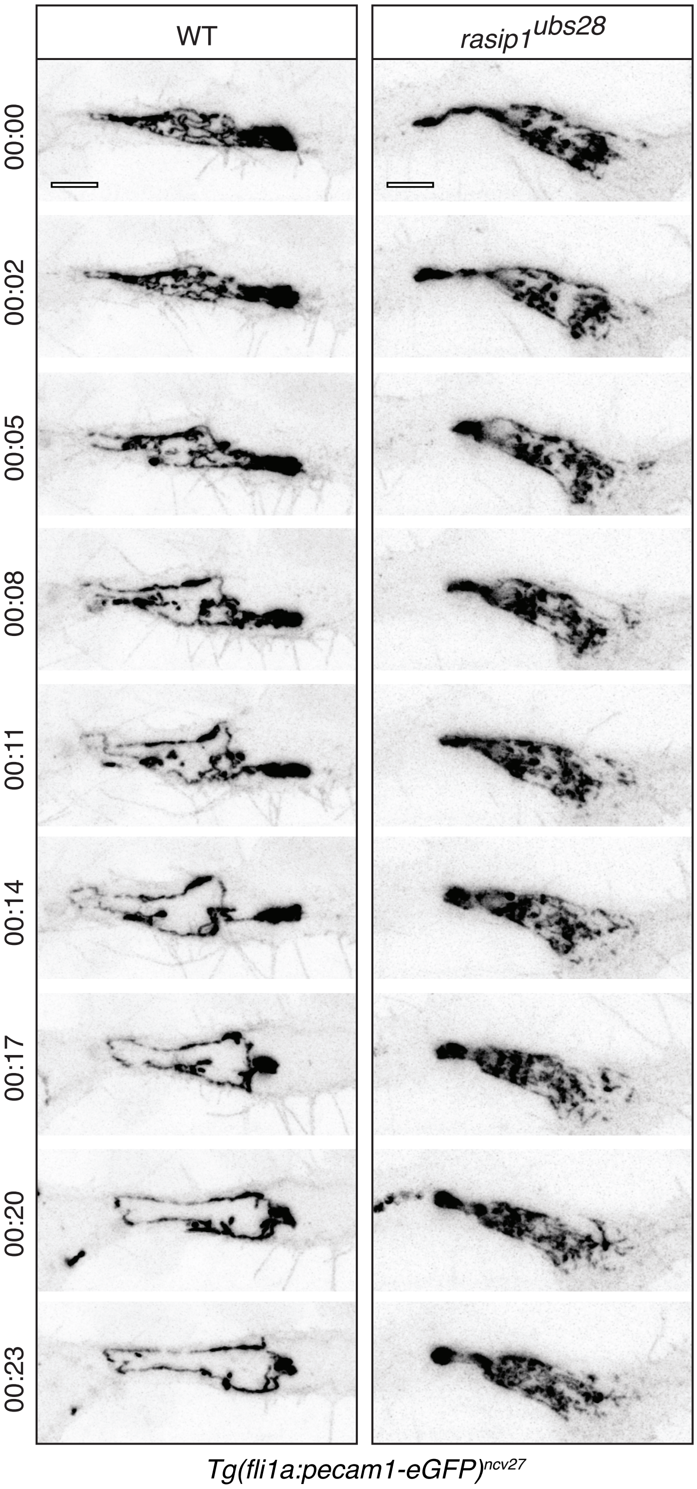


Supplementary Figure 3

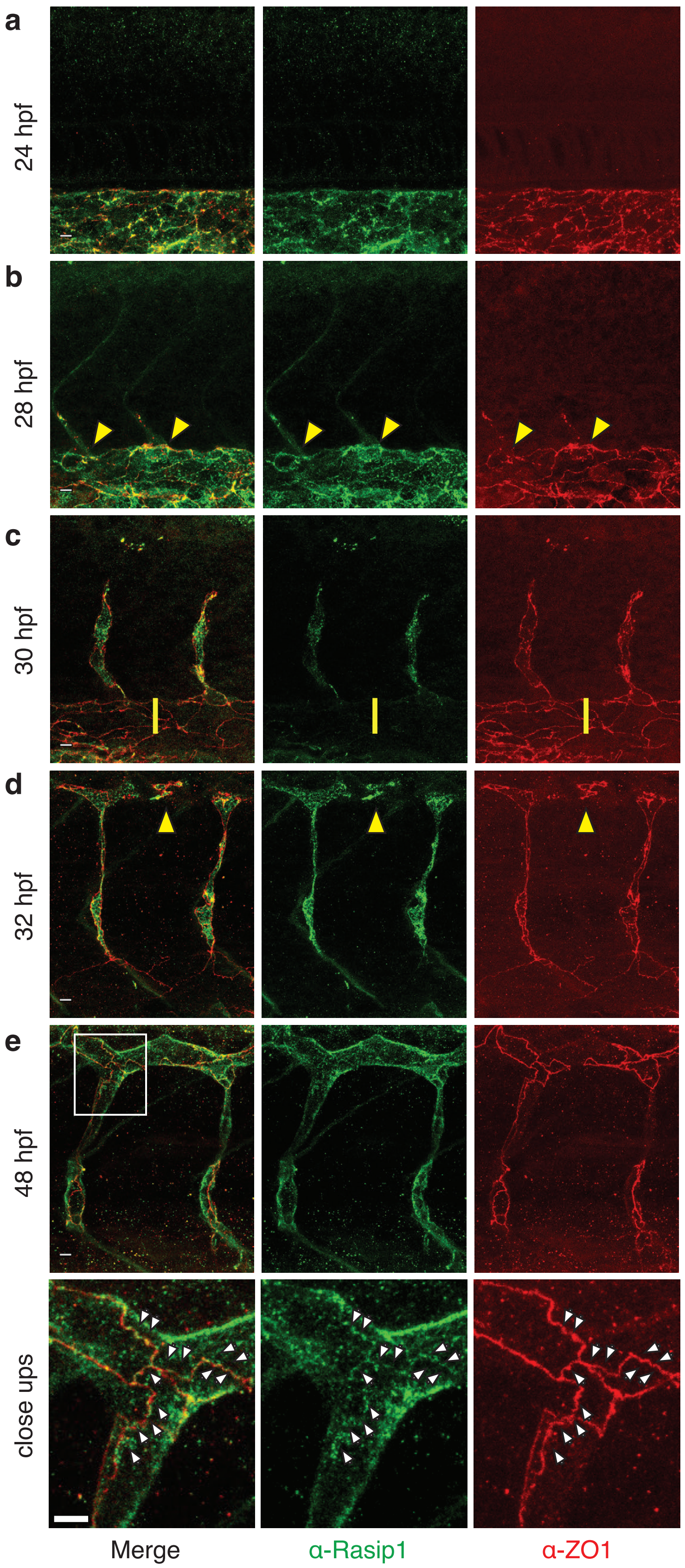


Supplementary Figure 4

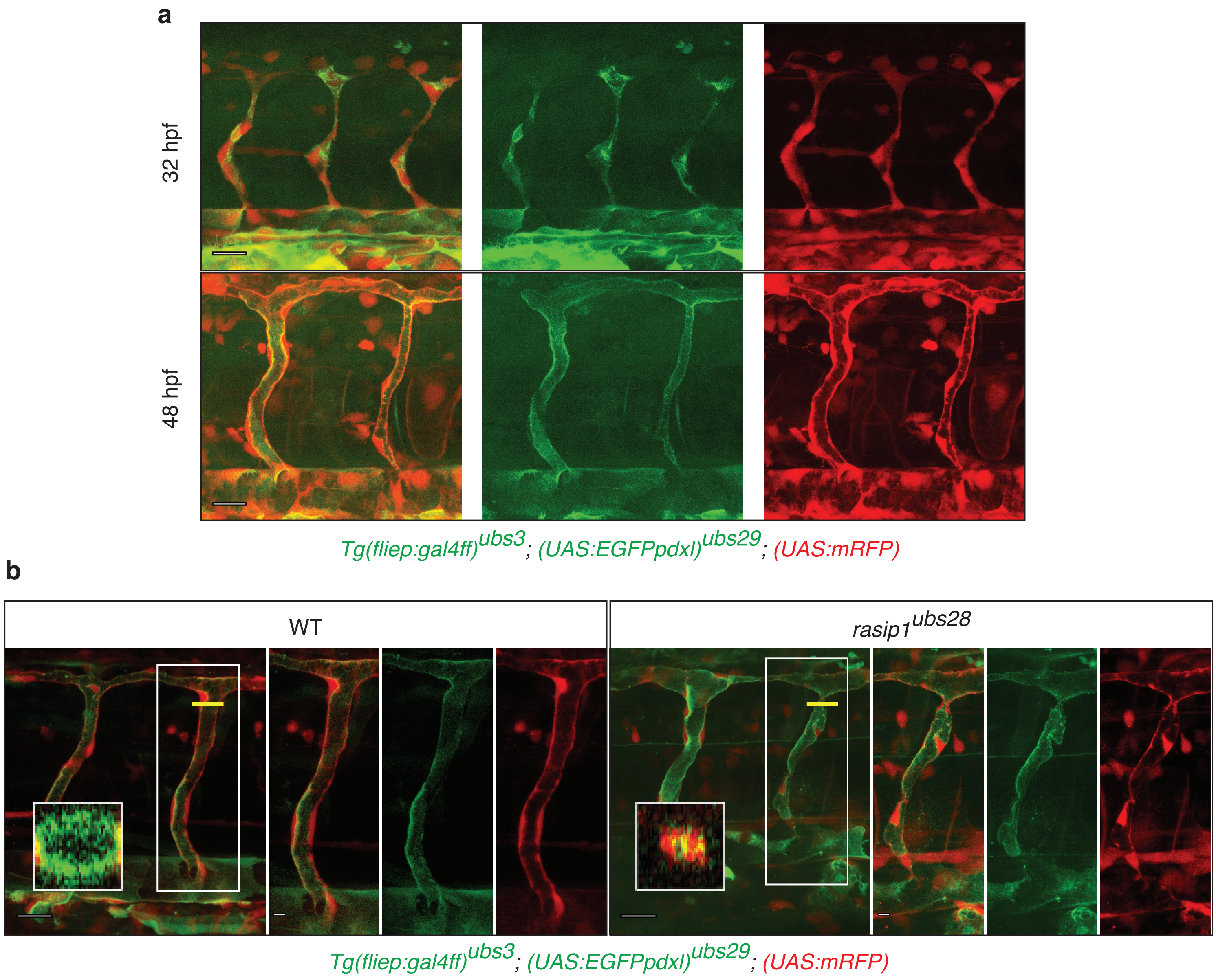


Supplementary Figure 5

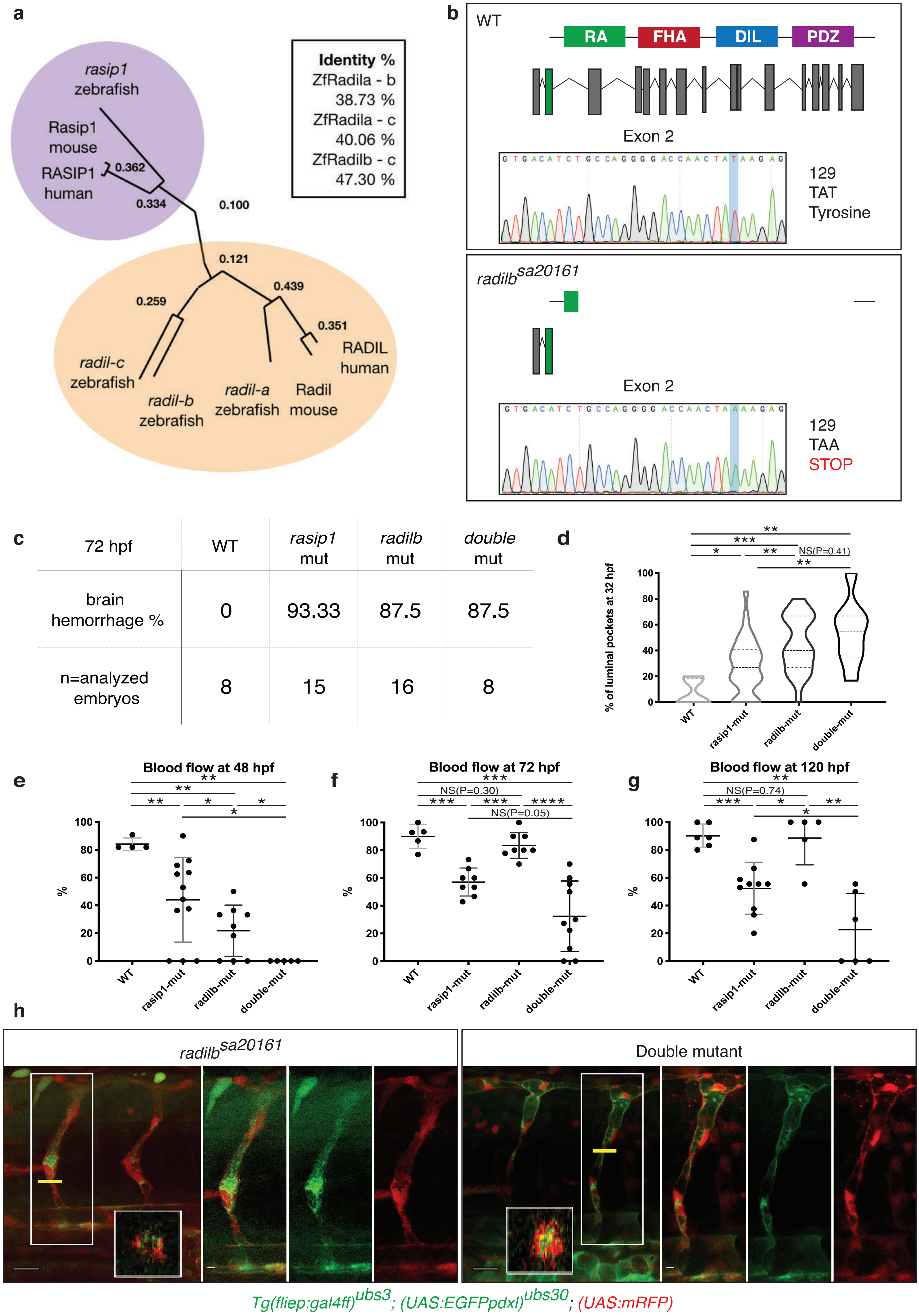


### Supplementary Figure 6

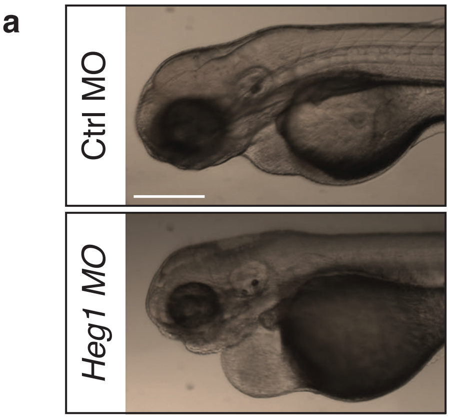


b

| 72 hpf | Ctrl MO | *Ccm1* MO | *Heg1* MO |
|---|---|---|---|
| brain hemorrhage % | 16.67 | 77.78 | 75 |
| n=analyzed embryos | 12 | 9 | 12 |

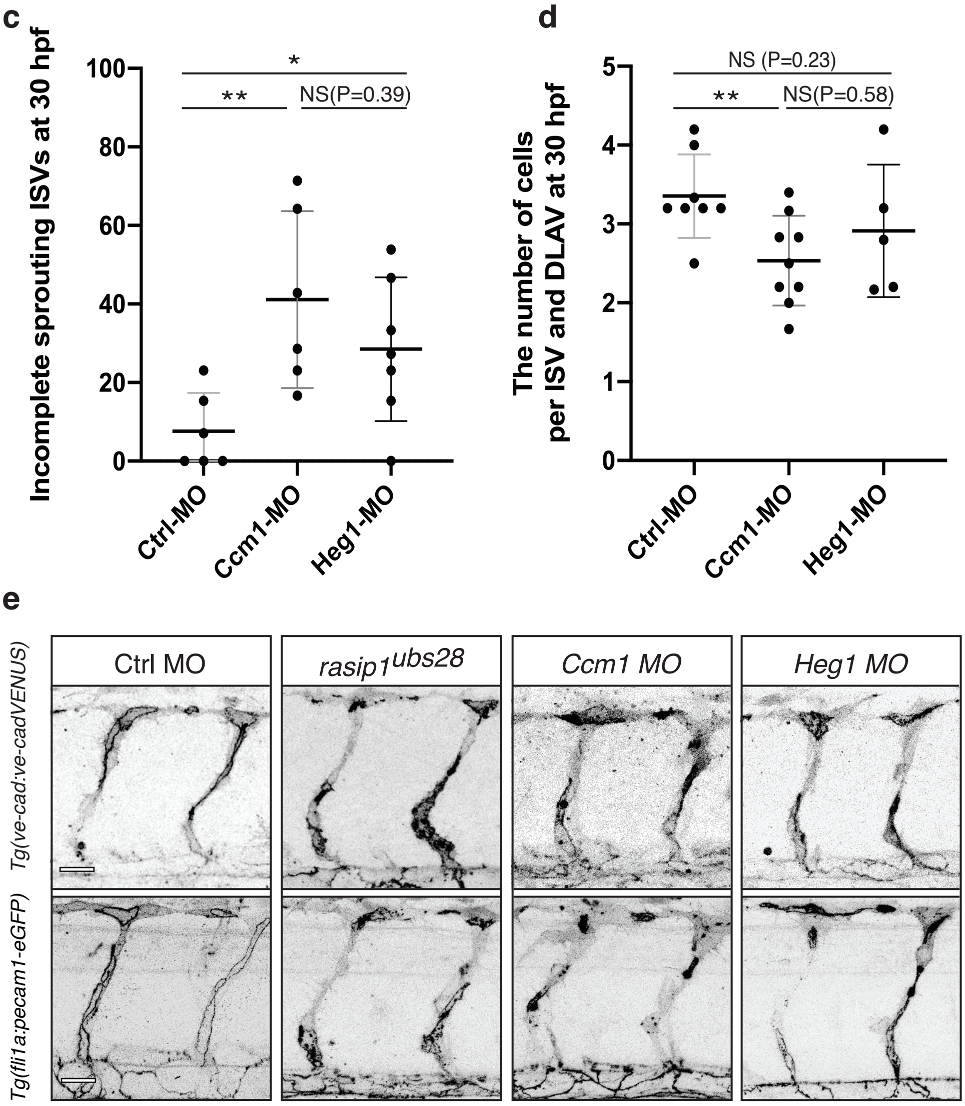


Supplementary Figure 7

### Supplemental Experimental Procedures

**Generation of *rasip1* mutant alleles**

For this study, three *rasip1* mutant alleles were generated. Two gRNA sites were selected for a null mutant (*rasip1$^{ubs28}$* mutant) using an online tool http://www.crisprscan.org (Moreno-Mateos et al., 2015) based on a high score:

Cris6 (GGCGGGGGAAGGGGATGGAGAGG, exon3, score 101) and

Cris7 (TGAAGCTCAGGGCTGGGGATTGG, exon16, score 63).

For *rasip1$^{ubs23}$* and *rasip1$^{ubs24}$* mutant, target sites were chosen:

Cris1 (GGAATGTCCCTTACAGCTGGTGG, exon3),

Cris2 (GGCGGGGGAAGGGGATGGAGAGG, exon 2),

Cris3 (GGACAAGACAGGTAGCGGAGGGG, exon12),

Cris4 (GGTGGAGTGAGAGAGGGAGG, exon2) and

Cris5 (GGCGGGACGGGAGTCACACGCGG, exon7).

gRNAs/Cas9 injections were performed according to (Gagnon et al., 2014). gRNAs were cloned into vector DR274. Injection mix: 1 µl Cas9 protein 6 mg/ml; 0.5 µl KCl 2M, 1 µl gRNA (around 1 µg/µl). Mutants were identified by sequencing the genomic target region.

**Genotyping of *rasip1* and *radil-b* mutant alleles**

*rasip1*and *radil* mutants were identified by multiplex-PCR using combinations of non-specific and allele-specific primers according to (Sauteur et al., 2014). Primer sequences are as follows:

| **Primer** | **Name** | **Sequence (5'-3')** |
|---|---|---|
| Rasip1-1 | Rasip1-fwd | TGTTGCCATCAGATCCACCAC |

| | | |
|---|---|---|
| Rasip1-2 | Rasip1-wt-rev | TTGGCCCGGGATTGCTGATT |
| Rasip1-3 | Rasip1-ubs28-rev | GTCCGCTGATTAGCAGGAAGT |
| Radilb-1 | Radil-b-fwd | CCACAACAACCGGCTAACCAC |
| Radilb-2 | Radil-b-rev | ACAATGAGCCTGGGTTGCAAATA A |
| Radilb-3 | Radil-b-wt-fwd | TGGCCAGCACACTCTTTT |
| Radilb-4 | Radil-b-sa20161-rev | GCCAGGGGACCAACTATA |

**Phylogenetic comparison of Rasip1 and Radil homologues**

The analysis was carried out using using the online program *phylo.io* (http://dev.phylo.io/#) (Robinson et al., 2016). The following peptide sequences were used: *Mus musculus* (mouse) Rasip1: ENSMUSG00000044562, *Homo sapiens* (human) Rasip1: ENSG00000105538, *Danio rerio* (zebrafish) Rasip1: ENSDART00000155407.3, *Mus musculus* (mouse) Radil: ENSMUSG00000029576, *Homo sapiens* (human) Radil: ENSG00000157927, *Danio rerio* (zebrafish) *radil-a*: ENSDARP00000101722, *radil-b*: ZDB-GENE-130530-682 si:ch73-281f12.4, *radil-c*: ZDB-GENE-121214-224 si:ch211-176g6.2.

### Supplementary Figure legends

**S-Figure 1: Characterization of *rasip1* mutants. (a, b)** Genomic organization of the *rasip1* locus in wild-type (a) and *rasip1*$^{ubs28}$ mutants (b). *rasip1* is encoded by 16 exons. gRNAs for CRISPR/Cas9 were designed to target exon3 and exon16. The wild-type DNA sequence of exon3 is shown. The *rasip1*$^{ubs28}$ mutant, which consists of a large deletion from exon 3 to exon 16, is lacking all exons encoding the three conserved protein domains. **(c)** PCR strategy to screen for *rasip1* mutant embryos or fish. **(d, e)** Schematic representation two additional mutant alleles, *rasip1*$^{ubs23}$ and *rasip1*$^{ubs24}$. **(f)** Immunofluorescen staining of Rasip1 (green) in the zebrafish vasculature (32hpf). The anti-zf-Rasip1 antibody is directed against the C-terminal domain of the protein (see Materials and Methods). The endothelium is labeled by *Tg(kdrl:EGFP)*$^{s843}$ (blue), junctions are labeled by Zo-1 (red). Rasip1 proteins is not detectable in *rasip1*$^{ubs28}$ mutants. Scale bars, 5 µm.

**S-Figure 2: ISV sprouting is affected in *rasip1* mutants. (a)** Still pictures of time-lapse movies (s-movies 1, 2) showing ISV sprouting in wild-type and *rasip1*$^{ubs28}$ embryos between 27 and 30 hpf. Scale bars, 20 µm. **(b)** Quantification of incomplete ISVs at 30 hpf (WT *n*=8 embryos, mut *n*=21). Median value: WT=0, mut=25%. Sprouting ISVs showing incomplete growth were counted and divided by the total ISV number per embryo. (unpaired two-tailed Mann Whitney test and error bars indicate standard deviation; significance: *p < 0.1) **(c)** Proportion of ISVs of different cell numbers at 30 hpf (WT *n*=12 embryos, 58 ISVs; mut *n*=12, 60). Unpaired two-tailed Mann Whitney test and error bars indicate standard deviation; significance: ***p < 0.001 **(d)** Cell division rates from 24 to 30 hpf are decreased in *rasip1*$^{ubs28}$ compared to wild-type. Embryos were analyzed by Fisher exact test: p=0.0316. **(e, f)** Still-pictures of time-lapse (s-movies 3-6) analysis showing endothelial cell proliferation and movements (visualized by nuclear EGFP) in wild-type and *rasip1*$^{ubs28}$ embryos. *rasip1* mutants show reduced cell proliferation within the sprout. Reduced cell number may be partially compensated by migration of additional cells into the sprout (f). Scale bars, 5 µm. **(g)** Quantification of time-lapse analyses on the relative contribution (%) of cell migration and proliferation to ISVs of different cell content. The ratio of cells from divisions and cells originated from the DA were quantified in *rasip1*$^{ubs28}$ compared to wild-type (WT *n*=4, mut *n*=5).

**S-Figure 3: Re-localization of Pecam1-EGFP from the apical region during anastomosis.** Still images from time-lapse movies (s-movies 12, 13) with high spatial and temporal resolution (hh:mm) from a movie of a PECAM-EGFP expressing embryo *Tg(fli1a:Pecam-EGFP)*$^{ncv27}$. Junctions were imaged in the DLAV from 32 hpf onwards. Scale bar, 5 µm.

**S-Figure 4: Dynamic distribution of Rasip1 during vascular development. (a-e)** Immunofluorescence of Rasip1 and Zo-1 in different developmental stages. Rasip1 protein is specifically expressed in the developing vasculature, visible in the DA at 24 hpf (a) and then in sprouting endothelial cells at 28 hpf (yellow arrowheads) (b). Expression in the DA is lost at 30 hpf (c). Rasip1 is apically localized at 30-32 hpf (yellow arrowheads) and also detectable at endothelial cell junctions at 48 hpf (white arrowheads in zoom-in). Scale bar, 20 µm.

**S-Figure 5: Loss of *rasip1* does not strongly affect apical polarization of endothelial cells. (a)** Live images showing localization of EGFP-Podocalyxin (EGFP-

Podxl) in the luminal cell membrane at 32 and 48 hpf. Scale bars, 20 µm. **(b)** Live images showing EGFP-Podxl in wild-type and *rasip1*$^{ubs28}$ *embryos. rasip1*$^{ubs28}$ mutants show local luminal constrictions (see inset z-projections). EGFP-Podxl is apically restricted in *rasip1*$^{ubs28}$ mutants but appears more irregular in its distribution. Scale bars, 20 µm (overview) and 5 µm (inset).

**S-Figure 6: Loss of *radil-b* enhances lumen defects and blood-flow of *rasip1* mutants. (a)** Phylogenetic tree based on the alignment of the entire protein sequences of human, mouse and zebrafish *rasip1* and *radil*. There are three *radil* paralogues in zebrafish. Numbers at branch points present bootstrap values. Zebrafish Radil-a has much closer relationship with regard to its mouse and human homologue based on protein-protein interaction databases. Radil-b and Radil-c were newly identified in this study and annotated from organism-specific databases. **(b)** A nonsense mutation in exon 2 of *radil-b*$^{sa20161}$ mutants ablates the Ras association (RA) domain, the dilute (DIL) domain (Rasip1 binding site) and the PDZ domain. **(c)** Quantification of cranial brain hemorrhage in *rasip1*$^{ubs28}$, *radil-b*$^{sa20161}$ and *rasip1*$^{ubs28}$; *radil-b*$^{sa20161}$ double mutants. **(d)** Quantification of luminal pockets from the still images of wild-type, single *rasip1*$^{ubs28}$ and *radilb*$^{sa20161}$ and *rasip1*$^{ubs28}$; *radilb*$^{sa20161}$ double mutants at 32 hpf. The number of ISVs containing ectopic lumens is divided by the total number of ISVs analyzed per embryo (WT *n*=5, *rasip1*$^{ubs28}$ mut *n*=34, *radilb*$^{sa20161}$ mut *n*=21, *rasip1*$^{ubs28}$; *radilb*$^{sa20161}$ mut *n*=8). **(e-g)** Quantification of blood flow defects in ISVs at 48, 72 and 120 hpf in *rasip1*$^{ubs28}$, *radilb*$^{sa20161}$ and in double mutants. *radilb*$^{sa20161}$ mutants show only transient defects in blood flow at 48 hpf. *rasip1*$^{ubs28}$;*radilb*$^{sa20161}$ double mutants show a strongly enhanced phenotype. Number of embryos and ISVs analyzed at 48, 72 and 120 hpf, respectively: WT (4, 44; 5, 71; 6, 87), *rasip1*$^{ubs28}$ (12, 176; 8, 119; 10,

118), *radilb*$^{sa20161}$ (9, 105; 8, 78; 5, 41) and double mutant (5, 56; 10, 102; 6, 55). Analyzed by unpaired two-tailed Mann Whitney test and error bars indicate standard deviation; significance (ns=no significance, *p < 0.1, **p < 0.01, ***p < 0.001, ****p < 0.0001). **(h)** Live images showing EGFP-Podxl in *radilb*$^{sa20161}$ and *radilb*$^{sa20161}$;*rasip1*$^{ubs28}$ double mutants displaying luminal constrictions at 48 hpf. Insets show digital cross sections of the ISV. Scale bars, 20 μm (overview) and 5 μm (inset).

**S-Figure 6: Vascular defects in *heg1* and *ccm1* morphants. (a)** At 72 hpf, *heg1* morphants show pericardial edema. Scale bar, 500 μm. **(b)** At 72 hpf, cranial brain hemorrhages are observed in *ccm1* and *heg1* MO injected embryos with higher incidence when compared to control MO injected embryos. **(c)** Quantification of incompletely sprouting ISVs at 30 hpf (control MO injected embryos *n*=6, *ccm1* MO *n*=6, *heg1* MO *n*=7). **(d)** The number of cells per ISV and DLAV at 30 hpf (Control MO injected embryos *n*=8, *ccm1* MO *n*=9, *heg1* MO *n*=5). **(e)** *In vivo* still images using VE-cad-VENUS and Pecam-EGFP as junctional reporters at 32 hpf. Scale bars, 20 μm. Analyzed by unpaired two-tailed Mann-Whitney test and error bars indicate standard deviation; significance (ns=no significance, *p < 0.1, **p < 0.01).

#### Supplementary Movie legends

**Supplementary Movie 1:** (S-Figure 2a) Confocal time-lapse movie of ISV formation (24-30 hpf) in wild-type embryos. Endothelial cells are labeled by TG(*(kdrl:EGFP)*$^{s843}$ (inversed contrast). Scale bar, 50 μm.

**Supplementary Movie 2:** (S-Figure 2a) Confocal time-lapse movie of ISV formation (24-30 hpf) in *rasip1*$^{ubs28}$ embryos. Endothelial cells are labeled by TG(*(kdrl:EGFP)*$^{s843}$ (inversed contrast). Compared to wild-type, *rasip1*$^{ubs28}$ mutants display unsynchronized and disrupted angiogenetic sprouting. Scale bar, 50 μm.

**Supplementary Movies 3-6:** (S-Figure 2e, f) Confocal time-lapse movie of ISV formation from 24 hpf in wild-type (s-mov3 and 5) and *rasip1*$^{ubs28}$ (s-mov4 and 6) embryos. Endothelial cells are labeled by *Tg(fliep:gal4ff)*$^{ubs3}$*; (UAS:mRFP)* in red and nuclei are labeled by *Tg(kdrl:EGFPnls)*$^{ubs1}$ in green. Scale bar, 20 μm.

**Supplementary Movie 7:** (Figure 3a) Confocal time-lapse movie of ISV formation (30-48hpf) in wild-type embryos. Endothelial cell junctions are labeled by VE-cad-Venus (*Tg(cdh5:cdh5-TFP-TENS-Venus)*$^{uq11bh}$) and imaged 1frame/h (reverse contrast). Scale bar, 50 μm.

**Supplementary Movie 8:** (Figure 3a) Confocal time-lapse movie of ISV formation (30-48hpf) in *rasip1*$^{ubs28}$ embryos. Endothelial cell junctions are labeled by VE-cad-Venus (*Tg(cdh5:cdh5-TFP-TENS-Venus)*$^{uq11bh}$) and imaged 1frame/h (reverse contrast). In *rasip1*$^{ubs28}$ mutants, collapsed junctions and VE-cadherin junction disconnections from the DA were observed. Scale bar, 50 μm.

**Supplementary Movies 9-11:** (Figure 3a-c) Confocal time-lapse movie of anastomotic ring formation in a wild-type(s-mov 9) and two *rasip1*$^{ubs28}$ mutant (s-mov 10, 11) embryos from 32hpf. Endothelial cell junctions are labeled by VE-cad-Venus (*Tg(cdh5:cdh5-TFP-TENS-Venus)*$^{uq11bh}$) (reverse contrast). Scale bar, 20 µm.

**Supplementary Movies 12 and 13:** (S-Figure 3) Confocal time-lapse movie showing dynamic re-localization of Pecam-EGFP (*Tg(fli1a:pecam1-eGFP)*$^{ncv27}$) during anastomosis in a wild-type (s-mov 12) and *rasip1*$^{ubs28}$ mutant (s-mov 13) embryo, starting at 32 hpf and recorded at 1 frame/min (00:00 to 00:43) In the *rasip1*$^{ubs28}$ mutant, a defect in the clearance of apical junctional proteins was observed. Scale bars, 5 µm.

**Supplementary Movies 14 and 15:** (Figure 4a) Confocal time-lapse movie showing lumen formation and the onset of blood circulation in a wild-type (s-mov 14) and *rasip1*$^{ubs28}$ mutant (s-mov 15) embryo starting at 32 hpf. Endothelial cells are labeled by EGFP (*Tg(kdrl:EGFP)*$^{s843}$*;* blood cell are labeled by DsRed *Tg(gata1:DsRed)*$^{sd2}$. In *rasip1*$^{ubs28}$ mutants blood circulation in the ISV and DLAV is delayed. Scale bars, 5 µm.

**Supplementary Movies 16 and 17:** (Figure 5a) Confocal time-lapse movie of ISV formation (24-30 hpf) in a wild-type (s-mov 16) and a *rasip1*$^{ubs28}$ mutant (s-mov 17) embryo. Endothelial cells are labeled by Tg(*(kdrl:EGFP)*$^{s843}$ (inversed contrast). Only the *rasip1*$^{ubs28}$ mutant shows formation of local lumens. Scale bar, 50 µm.

**Supplementary Movies 18-21:** (Figure 5d) Confocal time-lapse movie showing lumen formation in the ISV and DLAV from 34 hpf onwards in a wild-type embryo (s-mov 18 and 19) and *rasip1*$^{ubs28}$ mutant (s-mov 20 and 21) embryos. s-movies 18 and 20: Endothelial cells are labeled by cytoplasmic RFP (*Tg(fliep:gal4ff)*$^{ubs3}$*; (UAS:mRFP)*). Images were taken every 6 mins. (inverse contrast). Scale bars, 20 µm. s-movies 19 and 21: The same movie as s-movies 18 and 20, respectively showing merged channels: endothelial cells: red *(UAS:mRFP)*); endothelial cell junctions: green (VE-cad-Venus) ((*Tg(cdh5:cdh5-TFP-TENS-Venus)*$^{uq11bh}$)
